# Chemically induced proximity reveals mechanotransduction of a meiotic checkpoint at the nuclear envelope

**DOI:** 10.1101/2023.10.28.564506

**Authors:** Chenshu Liu, Abby F. Dernburg

## Abstract

Successful sexual reproduction relies on robust quality control during meiosis. Assembly of the synaptonemal complex between homologous chromosomes (synapsis) regulates meiotic recombination and is crucial for accurate chromosome segregation in most eukaryotes. Synapsis defects can trigger cell cycle delays and, in some cases, apoptosis. Here, by developing and deploying a new chemically induced proximity system, we identify key players in this quality control pathway in *Caenorhabditis elegans*. We find that persistence of the Polo-like kinase PLK-2 at pairing centers, specialized chromosome regions that interact with the nuclear envelope to promote homolog pairing and synapsis, induces apoptosis of oocytes by phosphorylating and destabilizing the nuclear lamina. Unexpectedly, we find that a mechanosensitive Piezo1/PEZO-1 channel localizes to the nuclear envelope and is required to transduce this signal to promote apoptosis. Thus, mechanosensitive ion channels play essential roles in detecting nuclear events and triggering apoptosis during gamete production.

**One-sentence summary:** Destabilization of the nuclear lamina triggers Piezo-dependent germline apoptosis.

## Introduction

A variety of surveillance mechanisms help to ensure accurate transmission of genetic information during mitosis and meiosis. In many eukaryotes, defects during meiosis or gametogenesis can lead to cell cycle delay and/or cell death. While some of these pathways respond to unrepaired DNA damage (1; 2), additional checkpoints monitor meiosis-specific events such as synapsis, defined as the assembly of the synaptonemal complex (SC) between homologous chromosomes (3–6). In some organisms, apoptosis is a more common outcome of defective meiosis in one sex than the other; for example, in mammals, defective spermatocytes are culled more stringently than oocytes (7; 8). This accounts for sterility (azoospermia) of men carrying mutations in genes required for meiosis (9; 10), and may also contribute to the higher incidence of aneuploidy in human ova than in sperm (8; 11).

In the hermaphroditic nematode *Caenorhabditis elegans*, meiotic defects can trigger cell cycle checkpoints in both spermatogenesis and oogenesis. Defective oocytes, but not spermatocytes, also frequently undergo apoptosis, even though both types of gametes are produced in the same tissue in hermaphrodites (12). A substantial fraction of oocytes (more than half) undergo “physiological” apoptosis even in the absence of apparent meiotic defects (13), but unrepaired DNA damage or failures in homologous chromosome synapsis result in elevated apoptosis and culling of the affected oocytes (3). Previous work has identified factors that are specifically required for apoptosis in response to DNA damage or synapsis defects. The former depends on canonical DNA damage sensing and signaling factors, while the “synapsis checkpoint” depends on functional pairing centers (PCs), special regions on each chromosome that mediate nuclear envelope attachment and promote homolog pairing and synapsis during early meiotic prophase (3; 14–17). It also requires the meiotic kinase PLK-2 and the widely-conserved AAA+ ATPase PCH-2 (3; 18). Yet how meiotic nuclei detect and respond to unsynapsed chromosomes remains unclear.

In early meiosis, the meiotic kinases CHK-2 and PLK-2 are recruited to chromosome PCs by zinc finger proteins that bind to dispersed motifs in these chromosome regions and interact with the nuclear envelope (14; 15; 19). These kinases phosphorylate SUN-1, a component of the Linker of the Nucleoskeleton and Cytoskeleton (LINC) complexes that promote chromosome movement along the nuclear envelope during early prophase (18; 20). PLK-2 also phosphorylates the lamin protein LMN-1, which makes the lamina more labile and facilitates chromosome movement along the nuclear surface (21). Upon completion of synapsis, PLK-2 relocalizes from PCs to the SC (18; 22; 23). Synapsis defects result in prolonged association of PLK-2 with PCs and extended phosphorylation of SUN-1 and LMN-1 (24– 26). PLK-2 is also required for the synapsis checkpoint (18). These findings led us to speculate that the persistence of PLK-2 at PCs might signal the presence of unsynapsed chromosomes.

## Results

### A chemically induced proximity system in *C. elegans*

To test whether PLK-2 at chromosome PCs affects meiotic progression or apoptosis, we adapted the auxin-inducible degradation (AID) system to control protein localization. In plants, the phytohormone auxin (indole acetic acid, IAA) triggers binding of target proteins carrying a “degron” sequence by the F-box protein TIR1, which interacts with Cullin1 and associated proteins to form an SCF E3 ubiquitin ligase complex (27). Expression of TIR1 from *Arabidopsis thaliana* in *C. elegans* results in auxin-induced degradation of proteins tagged with a degron sequence derived from Arabidopsis IAA17, or similar degron sequences (28; 29). We modified our *C. elegans* AtTIR1 expression sequence to incorporate 2 amino acid substitutions that abrogate the interaction between TIR1 and Cullin 1 and thereby block degradation of degron-tagged proteins (30; 31)(Figure 1A). We first expressed this mutated TIR1 (TIR1^CIP^) fused to mRuby, and confirmed that in the presence of auxin, this fluorescent fusion protein was recruited to proteins fused with a degron (e.g. PLK-2::AID) (Figure S1; Movie S1), but did not result in their degradation.

**Figure 1.**
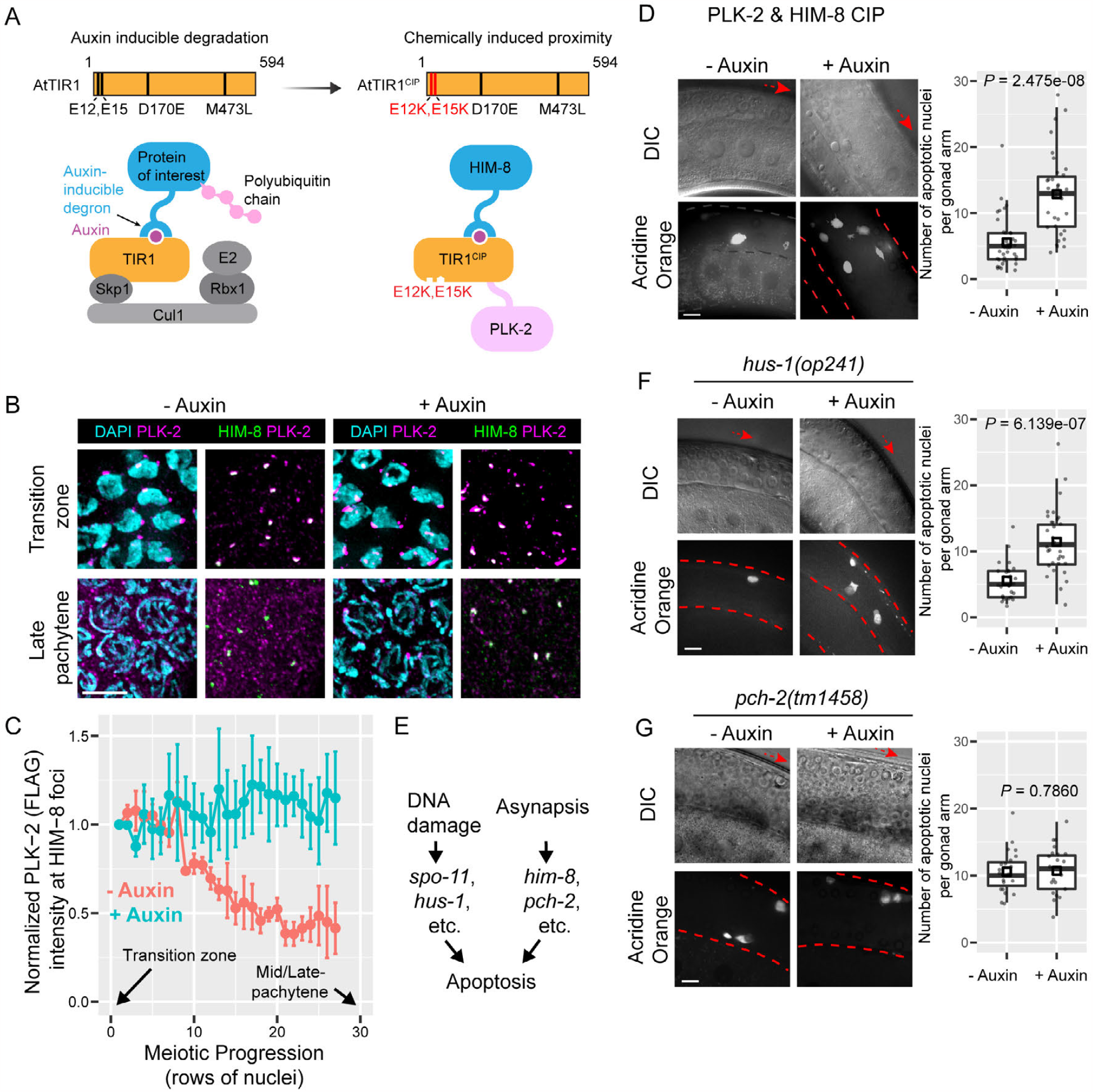
Recruitment of PLK-2 to X-chromosome pairing centers (PCs) activates the synapsis checkpoint. (A) The auxin-inducible degradation system was adapted to create an inducible dimerization system by incorporating 2 amino acid substitutions that abrogate binding between TIR1 and other E3 ligase components. (B) Immunofluorescence showing PLK-2 localization in relation to X-PCs, marked by HIM-8, in transition zone and late pachytene nuclei, with or without auxin treatment. Scale bar, 5 µm. (C) Quantification of PLK-2 (FLAG) intensity at HIM-8 foci during meiotic progression. Means ± SEM were calculated based on data from 3 animals. (D, F, G) Representative examples of DIC and acridine orange fluorescence images of germlines from the specified genotypes, ± auxin treatment. Scale bars, 10 µm. Boxplots show quantification of apoptotic nuclei based on AO staining. Each dot represents one animal. Medians (black crossbars) and means (black boxes) are shown. (E) Schema showing two meiotic checkpoints that lead to apoptosis.

This should be a broadly useful inducible dimerization/proximity system for *C. elegans*. Auxin is nontoxic, inexpensive, water-soluble and can readily penetrate the worm eggshell and cuticle at all stages of development, in contrast to larger inducers such as rapamycin (32–39). Many degron-tagged alleles of endogenous genes already exist and can be readily exploited for dimerization experiments. While auxin has some physiological effects on *C. elegans*, including a modest lifespan extension and ER stress resistance (40; 41), these will not generally affect the interpretation of experiments involving short-term manipulation of protein localization.

### PLK-2 at PCs is sufficient to induce apoptosis of oocytes

Pairing centers in *C. elegans* are chromosome regions required for homolog pairing and synapsis during meiosis. They are functionally defined by the presence of many binding sites that recruit one of 4 paralogous zinc finger (ZnF) proteins to each of the 6 chromosomes (14; 15). HIM-8 associates with the X-chromosome PC throughout meiotic prophase (15). We generated a strain expressing PLK-2::TIR1^CIP^ and degron-tagged HIM-8 (AID::HIM-8). In animals exposed to auxin, PLK-2 was detected at X chromosome pairing centers throughout meiotic prophase, while in control animals PLK-2 was only enriched at these sites in “transition zone” (leptotene/zygotene) and early pachytene nuclei (Figure 1, B and C) (22; 23). Importantly, induction of dimerization between PLK-2 and HIM-8 did not cause synapsis defects (Figure S2).

To quantify germline apoptosis, we stained live worms with the vital dye acridine orange (AO), which specifically labels apoptotic cell corpses in the germline (12; 42–46). Following imaging of intact animals using a spinning disk confocal microscope we counted the number of fluorescent corpses in each arm of the gonad. Ectopic targeting of PLK-2 to X-chromosome PCs significantly increased the number of apoptotic nuclei in the germline (Figure 1D). Elevated apoptosis was also detected when only one of the two *plk-2* genes was fused to TIR1^CIP^, demonstrating that PLK-2 recruitment to the X chromosome PC is sufficient to trigger oocyte death even when the untagged protein localizes to the SC (Figure S3). Either unrepaired DNA damage or unsynapsed chromosomes can induce elevated germline apoptosis (3) (Figure 1E). We found that HUS-1, an essential component of the DNA damage checkpoint, was dispensable for elevated apoptosis following recruitment of PLK-2 to X-PCs, while PCH-2, which is required for the synapsis checkpoint, was essential (Figure 1, F and G). Thus, ectopic targeting of PLK-2 to PCs triggers apoptosis via the synapsis checkpoint.

### Phosphorylation of the nuclear lamina promotes apoptosis

We next tested whether the kinase activity of PLK-2 is required to trigger apoptosis by introducing a previously-characterized “kinase-dead” mutation into the *plk-2::TIR1*^*CIP*^ fusion gene using CRISPR/Cas9 (21; 22) (Figure S4). Recruitment of PLK-2^K65M^ to HIM-8 did not induce germline apoptosis (Figure 2A), indicating that kinase activity is required for the role of PLK-2 in this checkpoint response.

**Figure 2.**
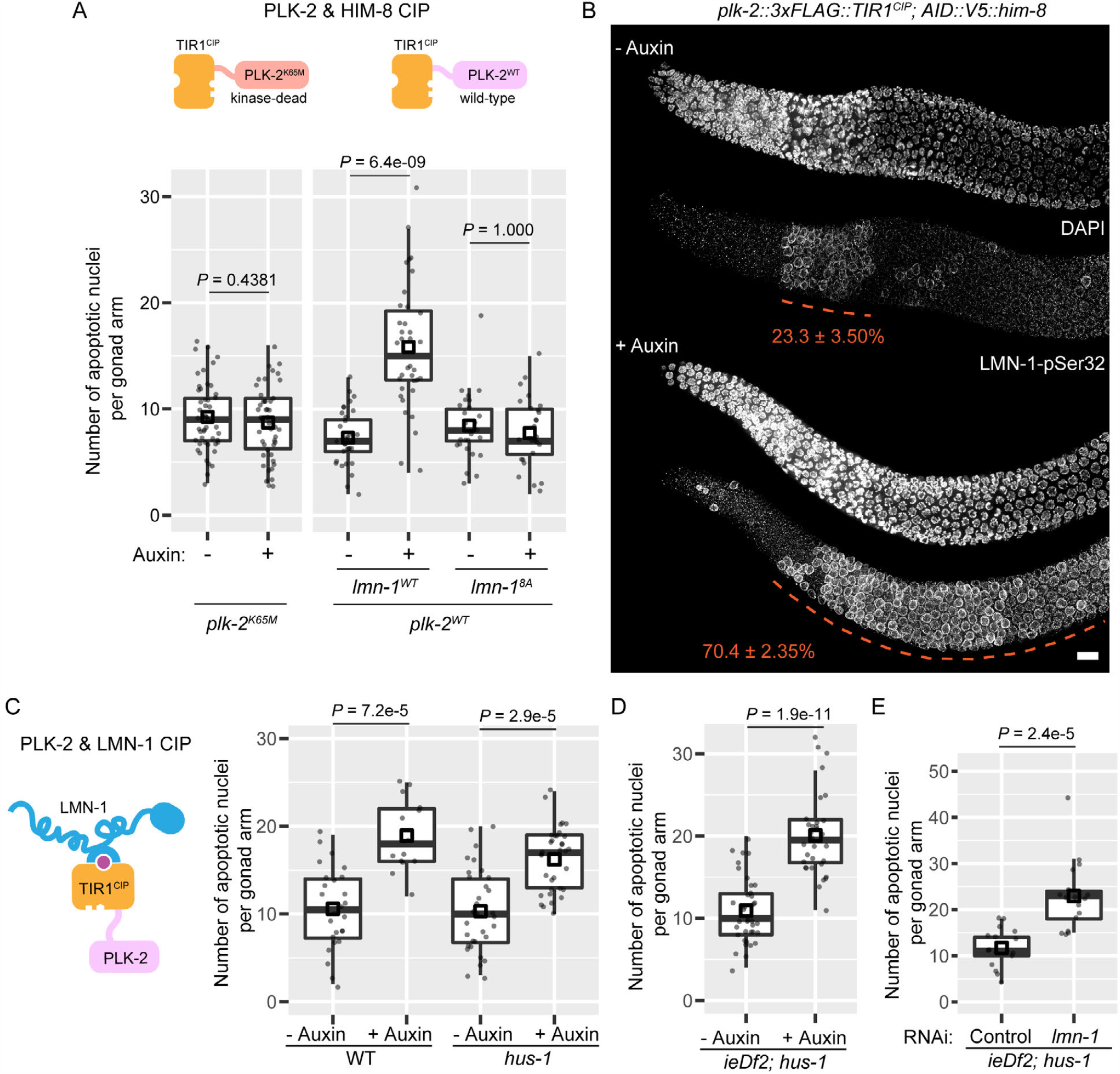
Phosphorylation of the nuclear lamina by PLK-2 promotes apoptosis. (A) Apoptosis in response to asynapsis requires PLK-2 kinase activity and LMN-1 phosphorylation. Recruitment of kinase-dead PLK-2^K65M^ to the X chromosome PCs fails to trigger elevated apoptosis. Mutation of PLK-2-dependent phosphorylation sites in LMN-1 also blocks induction of apoptosis. (B) Immunofluorescence showing extended region of meiotic nuclei with phosphorylated LMN-1-Ser32. Scale bar, 10 µm. Red dashed lines indicate regions with nuclei having phosphorylated LMN-1-Ser32. Numbers (means ± SD) indicate the percent of all meiotic prophase nuclei before late pachytene that stain positive for phosphorylated LMN-1-Ser32. *P* < 2.2e-16 by two-proportions z-test. (C) Recruiting PLK-2 to LMN-1 triggers apoptosis independent of HUS-1. (D) Recruitment of PLK-2 to LMN-1 is sufficient to trigger the synapsis checkpoint in the absence of functional PCs. *ieDf2* is a deletion of the 4 genes encoding essential PC proteins (18). *hus-1* is required for the DNA damage checkpoint. (E) Depletion of LMN-1 by RNAi also induces elevated apoptosis independent of HUS-1. For (A, C, D, E), Boxplots show quantification of apoptotic nuclei based on acridine orange staining. Each dot represents one animal. Medians (black crossbars) and means (black boxes) are shown.

Several serine and threonine residues in LMN-1 have been identified as PLK-2-dependent phosphorylation sites (18; 20; 21; 47). LMN-1 pSer32 is normally detected transiently following meiotic entry (21). Ectopic recruitment of PLK-2 to X-PCs greatly extended the region of meiotic nuclei positive for this marker (Figure 2B). We thus wondered whether lamin phosphorylation by PLK-2 is required for induced apoptosis. A mutant allele of LMN-1 in which 8 phosphorylation sites are mutated to alanine, LMN-1^8A^, reduces the solubility of the lamina in both mitosis and meiosis (21; 47). This allele prevented induction of germline apoptosis when PLK-2 was recruited to X-PCs (Figure 2A), indicating that LMN-1 phosphorylation is required for the checkpoint response.

We thus tested whether ectopic targeting of PLK-2 to the nuclear lamina is sufficient to induce germline apoptosis. Indeed, auxin-induced dimerization of PLK-2::TIR1^CIP^ and LMN-1::AID (45) resulted in elevated germline apoptosis that was independent of HUS-1 and DNA damage (Figure S5; Figure 2C). Moreover, by crossing in a deletion allele that disrupts all 4 of the PC zinc finger proteins, we found that recruitment of PLK-2 to the nuclear lamina bypassed the requirement for functional pairing centers in triggering apoptosis (Figure 2D). Thus, the crucial role of PCs in the synapsis checkpoint is to recruit PLK-2 to the nuclear envelope, where it phosphorylates LMN-1.

### Recruiting PLK-2 to LMN-1 weakens the nuclear lamina

We further investigated the effects of recruiting PLK-2 to the nuclear lamina through immunofluorescence. Upon PLK-2 recruitment, staining of epitope-tagged LMN-1 revealed a “dark zone” spanning much of meiotic prophase, indicating that LMN-1 at the nuclear envelope was reduced or easily extracted during tissue preparation (Figure 3A). This apparent weakening of the nuclear lamina was not observed when kinase-dead PLK-2 (PLK-2^K65M^) was recruited to LMN-1, indicating that it is a consequence of LMN-1 phosphorylation (Figure 3B).To determine whether this apparent weakening of the lamina was a potential cause or consequence of apoptosis, we introduced a mutation in Apaf-1/CED-4, which is required for germline apoptosis (3; 48; 49). In *ced-4* null mutant, ectopic targeting of PLK-2 to LMN-1 still resulted in a weakened nuclear lamina, as indicated by strongly reduced LMN-1 immunofluorescence (Figure 3C). This suggests that nuclear lamina weakening triggered by recruiting PLK-2 to LMN-1 is not downstream of germline apoptosis. HUS-1 was also dispensable for the induced LMN-1 “dark zone,” indicating that this response is independent of the DNA damage checkpoint (Figure 3D). Additionally, depleting LMN-1 even in the absence of functional PCs also induced HUS-1-independent apoptosis (Figure 2E), indicating that weakening of the nuclear lamina through LMN-1 phosphorylation or depletion is sufficient to activate the synapsis checkpoint.

**Figure 3.**
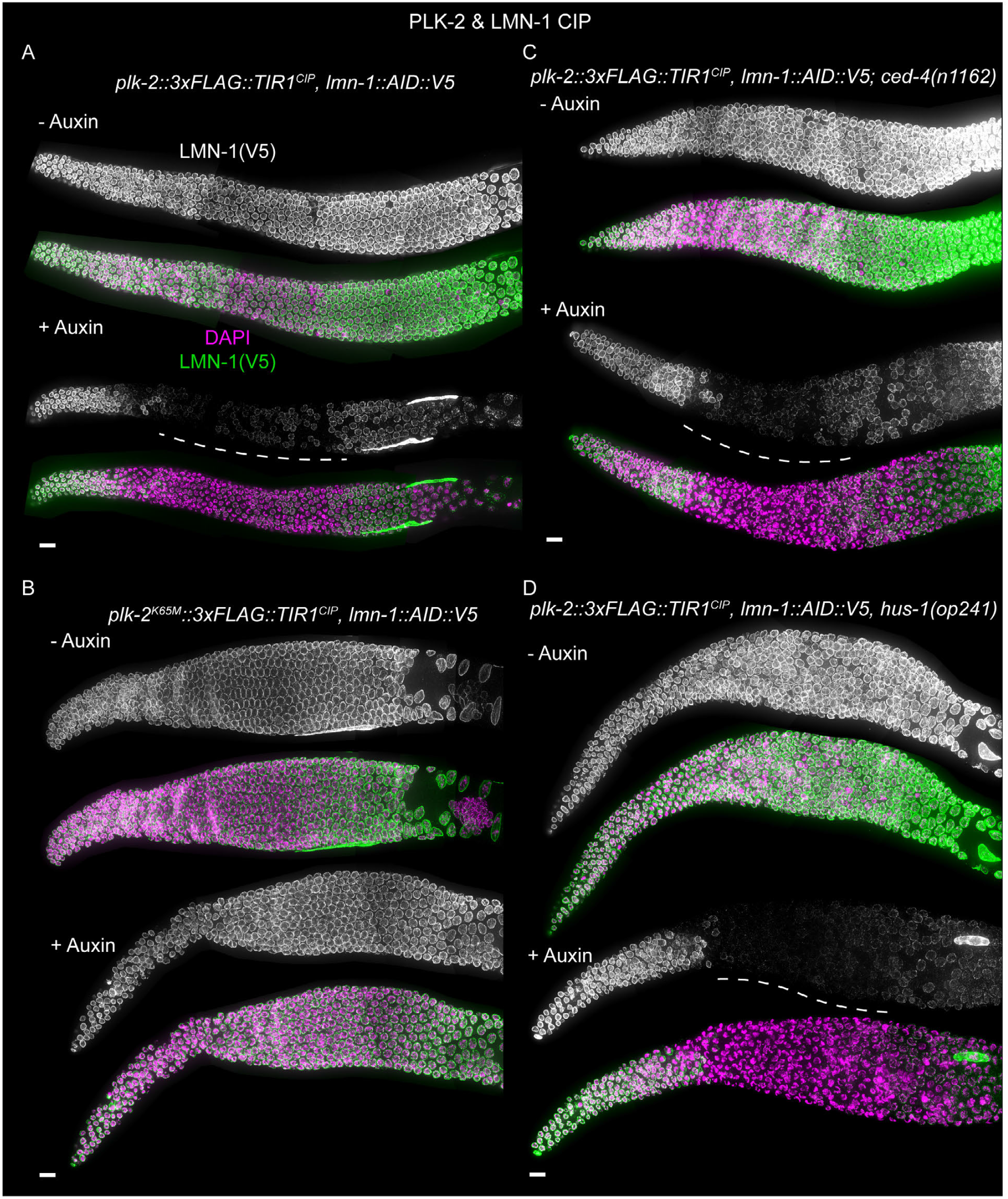
Recruitment of PLK-2 to LMN-1 weakens the nuclear lamina. Representative maximum-intensity projections of immunofluorescence images of gonads showing LMN-1::V5 (grayscale images and green in composite images) and DNA (DAPI, magenta in composite images). Scale bars, 10 µm. (A) to (D) each represents a different genotype. The lamin “dark zone” is indicated with dashed lines. LMN-1::V5 staining is still observed in the large, polyploid gonadal sheath cells that surround the germline cells.

**Figure 4.**
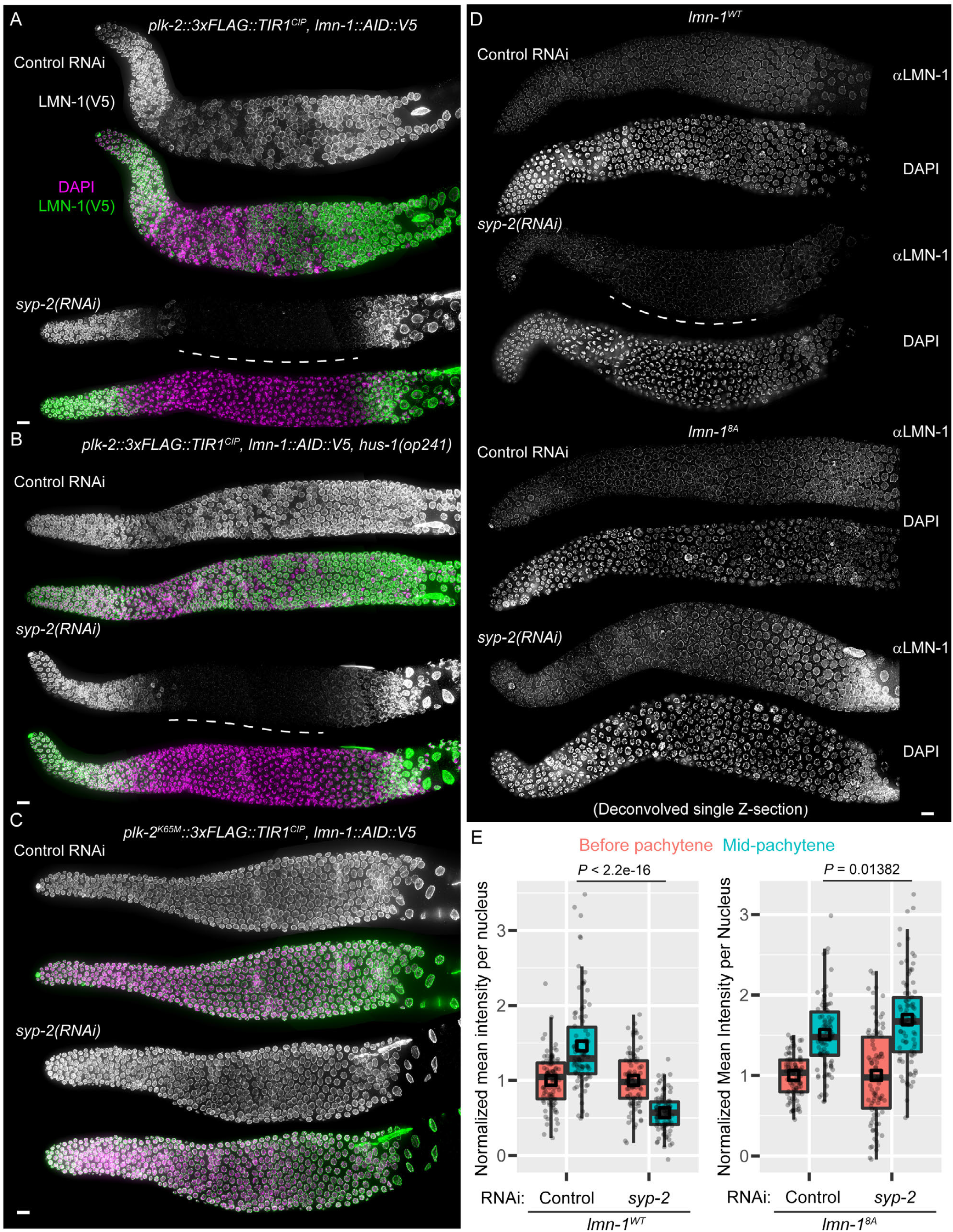
Asynapsis triggers PLK-2-dependent weakening of the nuclear lamina. (A) to (C) Representative images showing dissected gonads form animals treated with control RNAi or *syp-2(RNAi)*, stained for epitope (V5) tagged LMN-1 (grayscale images and green in composite images) and DNA (DAPI, magenta in composite images). Images are maximum-intensity projections. Scale bars, 10 µm. A lamin “dark zone” (dashed white lines) can be observed in wild-type (A) or in the absence of HUS-1 (B), but not in the absence of PLK-2 kinase activity (C). (D) Representative immunofluorescence of dissected gonads stained for endogenous LMN-1 in wild-type or *lmn-1*^*8A*^ background. Single optical sections from deconvolved data stacks are shown. Scale bar, 10 µm. The “dark zone” is more pronounced in animals expressing epitope-tagged LMN-1. (E) Boxplots show quantification of mean endogenous LMN-1 intensity per nucleus, normalized against endogenous LMN-1 intensity before pachytene. Each dot represents one nucleus. Medians (black crossbars) and means (black boxes) are shown. For each condition, 30 meiotic nuclei per stage from each of 3 animals were measured (Control RNAi or *syp-2(RNAi)*). *P* values were calculated using the two-sample t-test.

Mutation or depletion of essential SC components results in asynapsis and elevated germline apoptosis (Figure S6) (24; 50–52). We induced asynapsis by *syp-2(RNAi)* and found that this also resulted in an extended “dark zone” of LMN-1 (Figure 4A). This did not depend on HUS-1 (Figure 4B), but was abolished in the absence of PLK-2 activity (*plk-2*^*K65M*^) and in the *lmn-1*^*8A*^ mutant (Figure 4, C to E), indicating that PLK-2-dependent phosphorylation of LMN-1 is a normal consequence of asynapsis. Elevated germline apoptosis in response to SYP-2 depletion was suppressed by the LMN-1^8A^ mutation (Figure S7), further indicating that phosphorylation of the lamina by PLK-2 is integral to the synapsis check-point.

In wild-type animals, the pachytene region of the gonad typically displays some “straggler” nuclei with polarized chromosome morphology typical of the transition zone (19; 53; 54). These nuclei often have synapsis defects and also show a weakened nuclear lamina (Figure S8,A to D). We noted that the presence of straggler nuclei depends on the kinase activity of PLK-2 (Figure S8E; Figure 3; Figure 4, A and C).

Polarized nuclear morphology is seen following depletion of LMN-1 even in the absence of functional chromosome pairing centers (45). Together with the data presented here and observations from numerous other studies (19; 24; 25; 50; 52), this indicates that the primary driver of polarized, or “crescent-shaped” nuclear morphology normally observed during leptotene-zygotene is the weakening of the nuclear lamina through phosphorylation or lamin depletion, rather than NE attachment or movement of chromosomes.

### The mechanosensitive channel Piezo1/PEZO-1 is required for the synapsis checkpoint

We recently reported that depletion of nuclear lamins in the *C. elegans* germline results in reduced nuclear stiffness and higher deformability of the NE (45). During meiotic prophase, deformation of the nuclear surface is promoted by LINC complexes and cytoplasmic dynein (45; 55). We found that recruitment of PLK-2 to the nuclear lamina also resulted in increased deformation of the nuclear surface (Figure 5, A and B; Movie S2). Thus, phosphorylation of the lamina makes the NE more pliable and vulnerable to mechanical forces.

**Figure 5.**
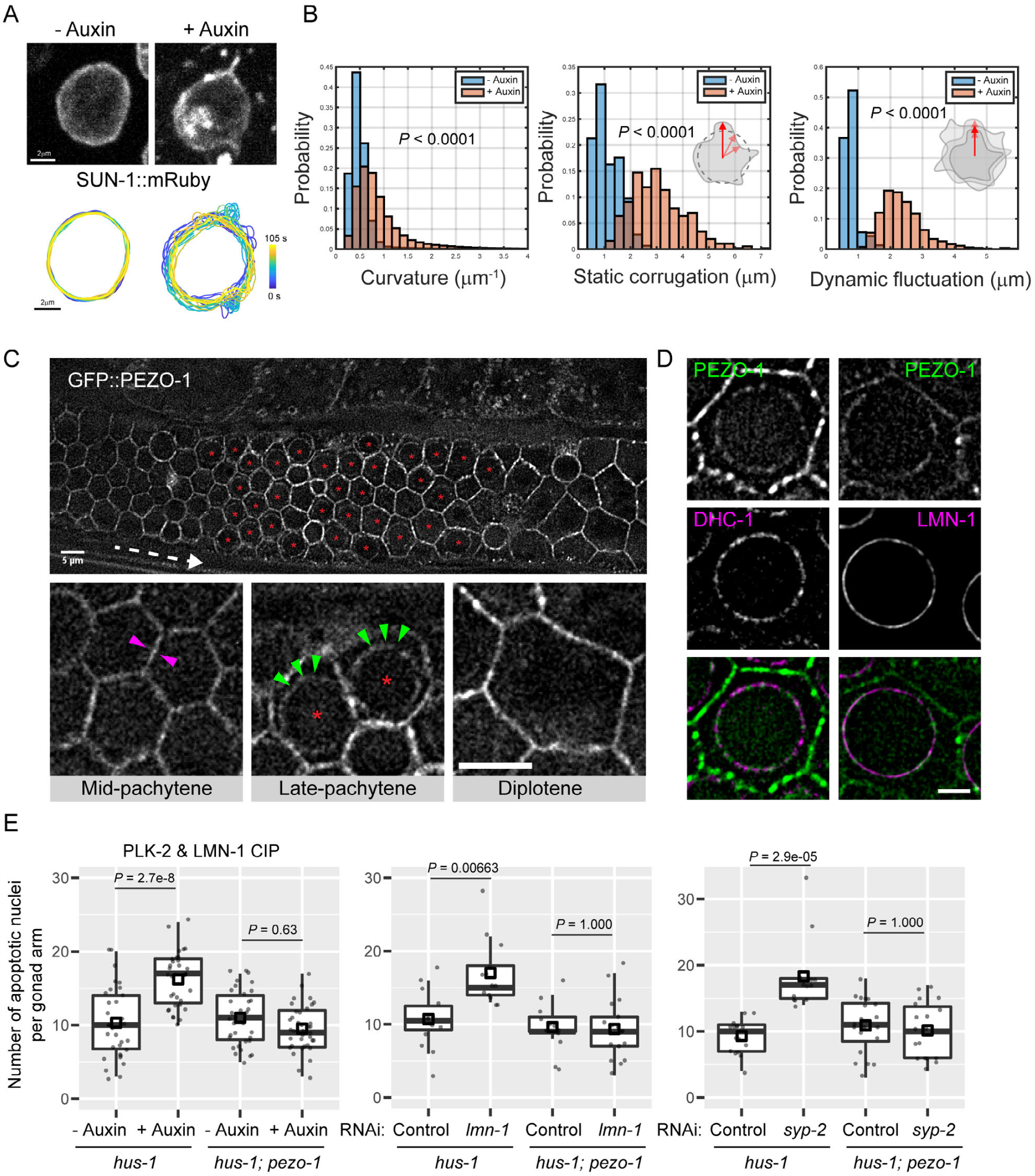
The mechanosensitive channel PEZO-1 localizes to the NE and is required for the synapsis checkpoint. (A) Still frames from live imaging of late pachytene nuclei expressing SUN-1::mRuby as an NE marker, without or with auxin treatment to recruit PLK-2 to the nuclear lamina. Images are maximum-intensity projections. Scale bar, 2 µm. Nuclear contours are aligned by centroids and color-coded by time. (B) Histograms for nuclear contours’ curvature, static corrugation and dynamic fluctuation. See Materials and Methods for more details. *P* values were computed with two sample t-test. (C) Live imaging of an adult gonad expressing GFP::PEZO-1 (single optical section). Dashed white arrow indicates the direction of meiotic progression. Red asterisks indicate meiotic cells with “ring-like” localization of GFP::PEZO-1 at the nuclear periphery. Magenta arrowheads indicate GFP fluorescence at the plasma membrane. Green arrowheads indicate GFP signals at the nuclear periphery. Scale bars, 5 µm. (D) Live imaging stills (single optical section) of late pachytene nuclei, showing localization of PEZO-1 and NE markers dynein (mCherry::DHC-1) or GFP::LMN-1 (21; 29; 45). GFP::PEZO-1 is in the left column and mScarlet::PEZO-1 in the right column. Scale bar, 2 µm. (E) PEZO-1 is required for CIP induced or asynapsis triggered apoptosis. Boxplots show quantification of apoptotic nuclei based on acridine orange staining. Each dot represents one animal. Medians (black crossbars) and means (black boxes) are shown.

Piezo channels are mechanosensitive ion channels that play key roles at the plasma membrane (56–59). *C. elegans* has only one Piezo channel, PEZO-1 (60; 61). GFP-tagged PEZO-1 localizes to plasma membranes throughout the *C. elegans* germline. Surprisingly, we also observed a population of PEZO-1 that appeared to be at the periphery of oocyte nuclei using two different fluorescently-tagged PEZO-1 strains (Figure 5C; Figure S9, A and B). We confirmed that a subset of PEZO-1 localizes at the NE (Figure 5D; Figure S9C), most prominently in the late pachytene region of the gonad, where apoptosis occurs (Figure 5C; Figure S9, A and B). We thus wondered whether PEZO-1 might be involved in detecting deformations of the nuclear envelope during oogenesis. In the absence of PEZO-1, ectopic recruitment of PLK-2 to the nuclear lamina did not lead to elevated germline apoptosis (Figure 5E). Moreover, recruiting PLK-2 to the nuclear lamina failed to trigger apoptosis when wild-type animals were exposed to Yoda1, a Piezo1 agonist that keeps the channel in an open state (60; 62). Yoda1 exposure had no effect on germline apoptosis in *pezo-1* mutants, indicating that it acts through PEZO-1. These results indicate that the ion channel function of PEZO-1 is required for the synapsis checkpoint (Figure S9D). Elevated germline apoptosis triggered by lamin depletion or asynapsis was also abolished in a *pezo-1* null mutant (Figure 5E). We further observed that the lamin “dark zone” caused by recruiting PLK-2 to LMN-1 does not require PEZO-1 (Figure S9E). Thus, the role of PEZO-1 in the synapsis checkpoint is downstream of lamin phosphorylation and destabilization.

### Asynapsis induces PCH-2-dependent redistribution of PEZO-1 at the NE

PEZO-1 localizes to both the plasma membrane and the NE, raising the question of whether the role of PEZO-1 in the synapsis checkpoint depends on channels within the nuclear membrane. We noticed that following SYP-2 depletion or recruitment of PLK-2 to the nuclear lamina, the distribution of GFP::PEZO-1 fluorescence at the NE changed markedly: in late pachytene nuclei a few foci were observed, rather than “ring-like” fluorescence along the NE. (Figure S10; Figure 6A). PEZO-1 foci colocalized with cytoplasmic dynein (DHC-1) and LINC complexes, which also clustered in oocytes affected by asynapsis (Figure 6B). This relocalization suggests that the pool of PEZO-1 at the NE is important for transducing a signal originating in the nucleus to trigger apoptosis. To our knowledge, Piezo channels have not previously been implicated in monitoring nuclear processes. SUN-1 and ZYG-12 constitute a LINC complex that transduces cytoskeletal forces to the NE throughout meiotic prophase (17; 45; 63). Depletion of SYP-2, which results in asynapsis, failed to trigger elevated apoptosis when SUN-1 was co-depleted, indicating that the presence of SUN-1 protein is also essential for the synapsis checkpoint (Figure 6C). A partial loss-of-function mutation, *sun-1(jf18)*, perturbs mechanotransduction by SUN-1/ZYG-12 LINC complexes, which results in nonhomologous synapsis and SPO-11-dependent apoptosis (64). In *sun-1(jf18)* homozygotes, SYP-2 depletion did not result in SPO-11-independent germline apoptosis (Figure 6C), indicating that LINC-mediated mechanotransduction is important for the synapsis checkpoint.

**Figure 6.**
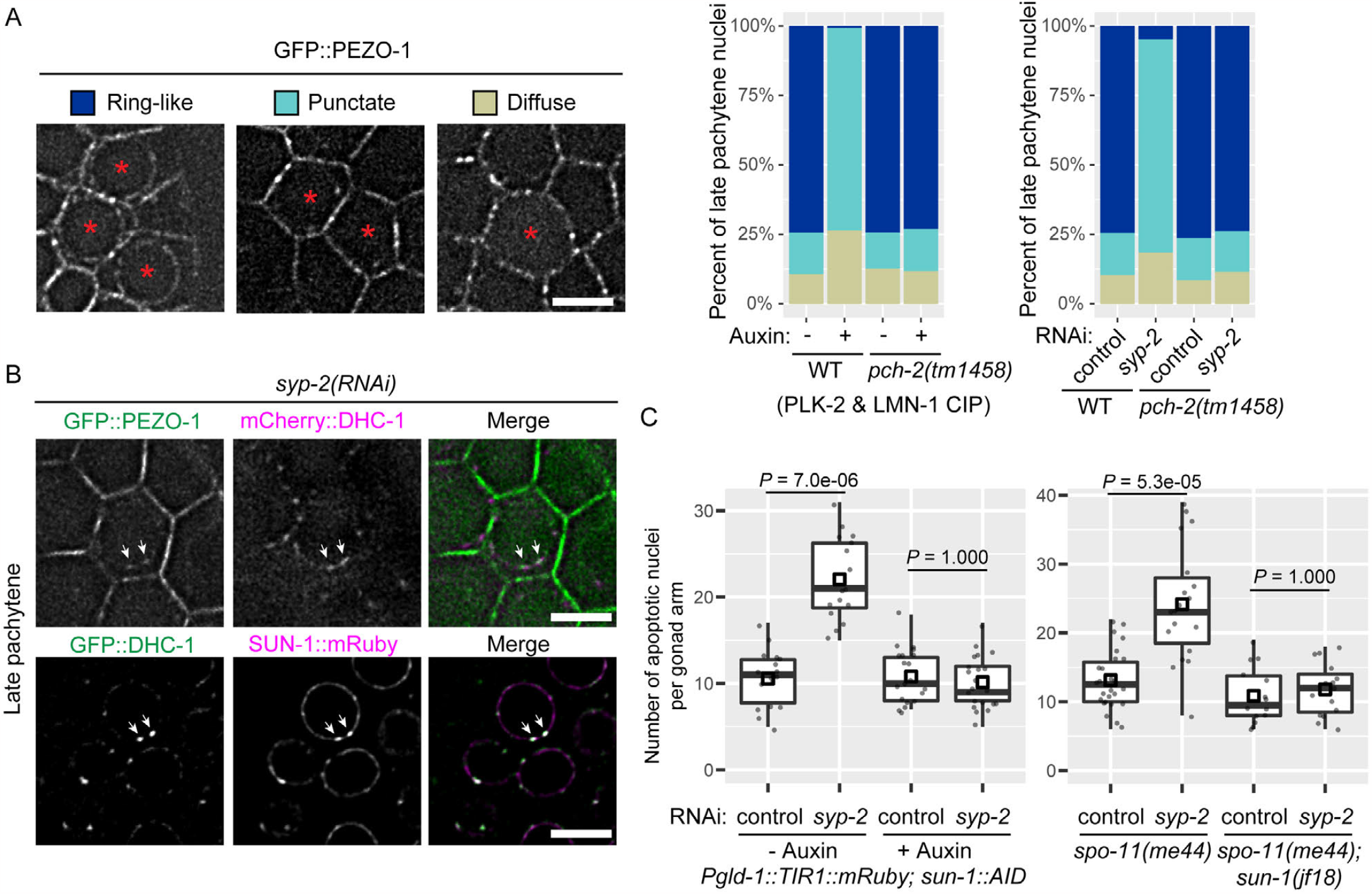
Asynapsis induces PCH-2-dependent PEZO-1 redistribution at the nuclear envelope. (A) Single confocal optical sections of images from live animals, illustrating three classes of GFP::PEZO-1 localization and their relative abundance among all late pachytene nuclei. Scale bar, 5 µm. Percentages were calculated from pooled data from at least three animals per condition. *P* < 2e-16 (WT – auxin vs. WT + auxin), *P* = 0.98 (*pch-2* – auxin vs. *pch-2* + auxin), *P* < 2e-16 (WT; control vs. WT; *syp-2(RNAi)*), *P* = 0.88 (*pch-2*; control vs. *pch-2*; *syp-2(RNAi)*), all computed using pairwise comparison of proportions (adjusted by the Benjamini-Hochberg method). (B) Representative images of live animals showing PEZO-1 foci that overlap with dynein (mCherry::DHC-1) and LINC complexes (SUN-1::mRuby) following *syp-2(RNAi)*-induced asynapsis. Images are single confocal optical sections of late pachytene nuclei in live animals. Scale bars, 5 µm. (C) Mechanotransduction mediated by SUN-1 is required for the synapsis checkpoint. Boxplots show quantification of apoptotic nuclei based on AO staining. Each dot represents one animal. Medians (black crossbars) and means (black boxes) are shown.

The AAA+ ATPase Pch2/PCH-2 plays a conserved but enigmatic role in meiotic checkpoints (65–72). Disruption of *pch-2* did not perturb the retention of PLK-2 at PCs or lamin phosphorylation (Figure S11; Figure S12); however, the redistribution of PEZO-1 at the NE in response to asynapsis was abrogated (Figure S10; Figure 6A). Thus, PCH-2 is required for the redistribution or remodeling of PEZO-1 at the NE. These observations further support a role for PEZO-1 at the NE in transducing apoptosis.

## Discussion

Chemically induced proximity (CIP) is a versatile experimental tool that takes advantage of cell-permeant small molecules (73). Several CIP approaches have been engineered for protein dimerization in cultured cells (31; 74), but their application in model organisms has been more limited, due to the challenges of delivering the ligands and their impact on physiology. By repurposing the versatile auxin-inducible degradation system, we developed a CIP system that is easy to use and can also capitalize on the plethora of *C. elegans* strains expressing degron-tagged endogenous proteins. This approach can enable (a) inducible visualization of endogenous proteins that are sensitive to constitutive fluorescent protein tagging in live animals (e.g. using fluorescently tagged TIR1^CIP^), and (b) ectopic recruitment and relocalization of proteins of interest. Here, the system has revealed a new quality control mechanism at the nuclear envelope during meiosis, where unsynapsed chromosomes lead to destabilization of the NE and Piezo/PEZO-1 dependent apoptosis (Figure 7). Cell nuclei experience mechanical forces from both intracellular and extracellular sources (75–78). Piezo1 has previously been implicated in transducing extra-cellular force to mediate changes in heterochromatin stiffness within the nucleus (79). This pathway involves Piezo1-mediated release of intracellular calcium from the endoplasmic reticulum (ER) and leads to a reduction in lamina-associated H3K9me3 heterochromatin, resulting in nuclear softening. Patch-clamp experiments on isolated nuclei have detected stretch-activated Ca^2+^ currents (80–82), suggesting the presence of mechanosensitive channels at the NE, but these were never identified. Our findings reveal that Piezo channels can also respond to forces acting directly on nuclei and lead to a cellular response. Based on known modes of activation for Piezo channels (56; 83; 84), we speculate that direct coupling between PEZO-1 and LINC complexes or sudden change of the local NE curvature open the Piezo channels at the nuclear membrane. This may lead to Ca^2+^ release from the perinuclear space or the ER lumen, triggering downstream pro-apoptotic pathways (Figure 7) (85–87).

**Figure 7.**
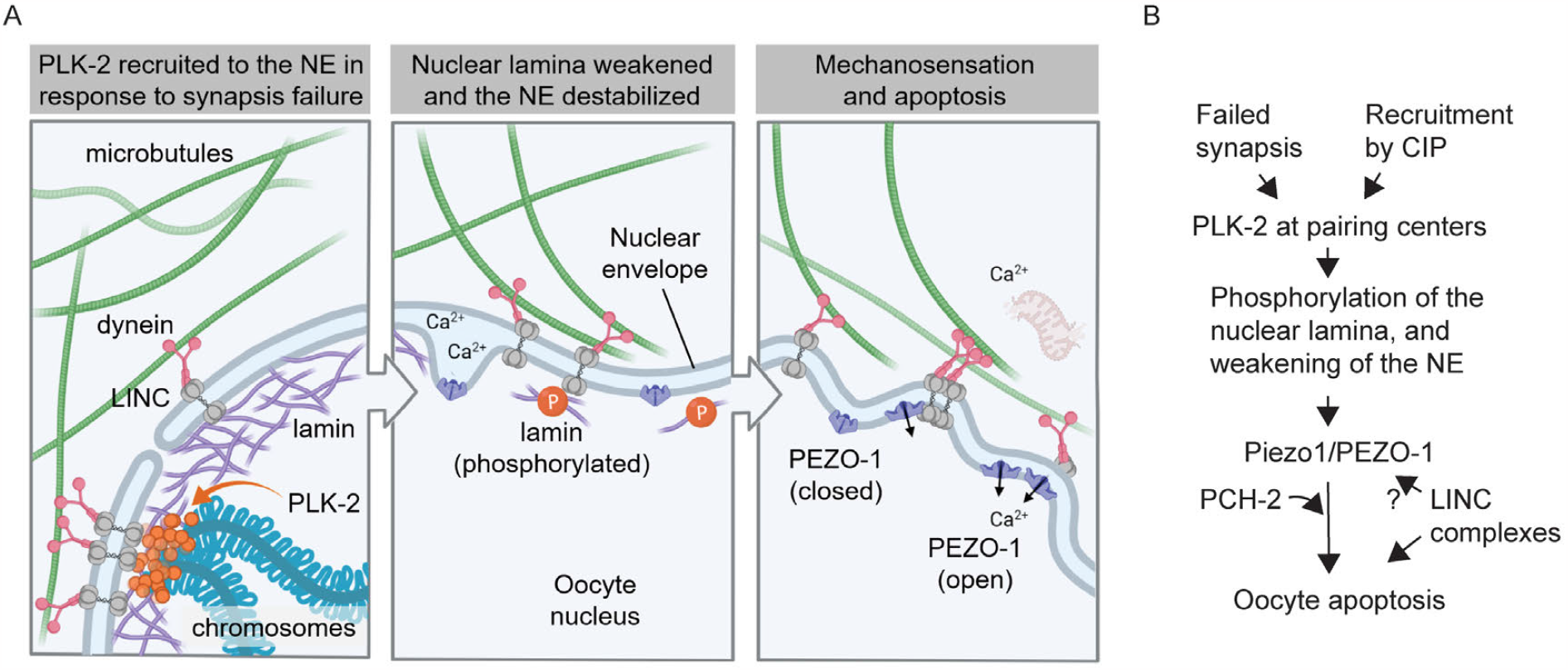
Model of a mechanosensitive checkpoint mechanism that monitors oocyte quality at the nuclear envelope. **(A)** In response to synapsis defects, PLK-2 is retained at the NE by binding to PCs. Phosphorylation of LMN-2 causes NE destabilization and PEZO-1-dependent apoptosis. (B) Schematic summary of the checkpoint pathway.

In most eukaryotes, telomeres interact with LINC complexes during early meiotic prophase and drive chromosomal motion that promotes homolog pairing (88). This evolutionarily conserved function can also be mediated by centromeres, as in *Drosophila* (89), or pairing centers, in *C. elegans* (88). Our findings reveal that these interactions between chromosomes and the nuclear envelope play an additional role in triggering a meiotic checkpoint via lamin phosphorylation. In *C. elegans*, homologous synapsis occurs independently of DNA double strand breaks (DSBs) or crossover (CO) formation. However, in most organisms that have been studied, SC assembly is triggered by the formation of crossover precursors. Monitoring of SC assembly thus serves as a crossover assurance mechanism. We suggest that a role for interactions between chromosomes and the NE in checkpoint signaling may also be widely conserved. In mice, the kinase Cdk2 localizes to both telomeres and CO sites, and is essential to link telomeres to the NE to form the “meiotic bouquet” (90; 91). Vertebrate CDKs phosphorylate lamins to disassemble the NE in mitosis (92–95) and Cdk2 may thus also modulate NE dynamics during meiotic prophase. DSB formation, which is essential for CO formation, is important for timely bouquet resolution (96). The bouquet stage is also prolonged in *Sycp3* mutants, which fail to complete synapsis (97). Additionally, mouse spermatocytes lacking the meiosis-specific lamin C2 show highly elevated apoptosis (98). While this may be a consequence of meiotic defects arising from the absence of meiotic lamins, it is also possible that perturbation of the lamina is a trigger for apoptosis, as we have demonstrated in *C. elegans*.

Spontaneous, or “physiological,” apoptosis of *C. elegans* oocytes results from a “hydraulic instability” that amplifies oocyte volume differences during a period of rapid cell growth, causing some cells to grow and survive while others shrink, leading to cell death (99). PEZO-1-mediated signaling may thus trigger apoptosis by predisposing affected cells to shrink and die during oocyte maturation.

In other organisms, oocytes also experience a range of mechanical forces during meiosis and oogenesis (45). In mammals, oocytes are produced in the ovary of a developing fetus and then arrest at the dictyate (diplotene) stage of meiosis for as long as several decades (100). Mechanical forces have been implicated in maintenance of this long arrest, and are likely to contribute to oocyte maturation and follicle selection as well as maintenance of ovarian reserve (101– 103). Piezo2 is detected in human ovarian follicles, (https://www.proteinatlas.org/ENSG00000154864-PIEZO2/tissue/ovary#img, from v23.0.proteinatlas.org), suggesting a potential role for these channels in mechanical regulation of dormancy and maturation (104). Our work reveals an important role for intracellular forces in oocyte quality control, with potential implications for age-related decline in oocyte quality and *in vitro* gametogenesis.

## Materials and Methods

### Generation of worm strains

All *C. elegans* strains were maintained at 20°C under standard conditions (105). All alleles were generated using CRISPR/Cas9 genome editing (Alt-R CRISPR-Cas9 gRNA products, IDT), as previously described (45). Details of alleles generated in this study are listed in Table S1. A complete list of *C. elegans* strains used and generated in this study is also provided in Table S2. Custom designed crRNAs targeting genes of interest were purchased from IDT. These sequences, as well as the sequences of dsDNA repair templates (gBlock from IDT) and ssDNA repair template for *dpy-10* (IDT) can be found in Table S3. A repair template for editing endogenous *plk-2* gene to express PLK-2::3xFLAG::TIR1^CIP^ was assembled into a pCR-Blunt backbone using Gibson Assembly (NEB) and confirmed using Sanger sequencing and whole-plasmid Nanopore sequencing (Primordium). Templates and crRNAs to create additional transgenes or alleles were designed and injected, but resulted in gene silencing, including the insertion of *TIR1*^*CIP*^*::V5* at the 5’ end of the endogenous *him-8* gene, insertion of a *TIR1*^*CIP*^*::V5::him8* transgene into the universal MosSCI site oxTI179 II, and replacing *mRuby* in *ieSi38[sun-1p::TIR1::mRuby::sun-1 3’UTR, Cbr-unc-119(+)] IV* with *V5::him-8*.

### Acridine Orange staining

Quantification of apoptosis by Acridine Orange (AO) staining was carried out as previously described (43; 45) with minor modifications. AO staining solution was prepared fresh by diluting a 10 mg/ml stock solution (Invitrogen™ A-3568) 1:200 into M9 buffer. Age-matched adult hermaphrodites were picked to the middle of a lawn of E. coli OP50 on fresh plates. 2 mL of staining solution was gently pipetted onto each plate until the entire surface (including the OP50 lawn) was covered. When necessary, animals were repositioned using a pipette onto the OP50 lawn to ensure active feeding behavior and efficient AO uptake. Plates were incubated for 1 hour at room temperature in the dark. Worms were then washed off the plate with M9 buffer, transferred to low-retention 1.5mL tubes (Fisherbrand) and pelleted by a low-speed spin (10 sec) in a tabletop mini centrifuge. They were washed with 600 µL M9 buffer. Pelleting and washing was repeated for a total of 3 washes, after which worms were transferred to fresh plates to recover at room temperature for 45 minutes in the dark. They were mounted on agarose pads and imaged within 1 hour of the end of the recovery period. Low retention pipette tips (Fisherbrand) were used throughout the washing process.

### RNA interference (RNAi), auxin and Yoda1 treatment, and apoptosis assays

Depletion of gene products by feeding RNAi was carried out as previously described (45). Bacterial strains from the Ahringer RNAi feeding library (106; 107) carrying RNAi clones targeting *syp-2* and *lmn-1* were validated by whole-plasmid Nanopore sequencing (Primordium) (see Table S4).

NGM worm plates containing 2mm indole acetic acid (IAA, auxin) were prepared and used as previously described (45). For experiments using the Piezo agonist Yoda1 (Tocris 5586), the compound was dissolved in DMSO (Amresco WN182) to make a 2.5 mM stock. This solution was diluted 1:125 into NGM medium to make plates cointaining 20 µM Yoda1(60). An equal volume of DMSO was added to control plates (60). For all experiments involving RNAi, auxin or Yoda1 treatment, and/or acridine orange staining, apoptosis was scored within 36 hours after the L4 stage to minimize any age-related effects on apoptosis (108).

### Immunofluorescence

Immunofluorescence was carried out as previously described (45; 109). The following antibodies were used: anti-FLAG (mouse M2, F3165, 1:400), anti-V5 (mouse, P/N 46-0705, Invitrogen, 1:400), anti-V5 (rabbit, 1:400), anti-GFP (mouse monoclonal, Roche, 1:400), anti-SYP-1 (goat, affinity purified, 1:400 (18)), anti-SYP-2 (rabbit, affinity purified, 1:1000), anti-HTP-3 (chicken, 1:400 (110)), anti-HTP-3 (guinea pig, 1:400 (110)), anti-HIM-8 (rat, 1:400 (15)), anti-HIM-8 (rabbit polyclonal, affinity purified, SDIX, SDQ2975, 1:1000), anti-NPP-7 (rabbit polyclonal, affinity purified, SDIX, SDQ0870, 1:1,000), anti-LMN-1 (rat, 3933, 1:200, a gift from Yosef Gruenbaum), anti-LMN-1-pSer32 (1:10 with pre-adsorption, a gift from Verena Jantsch (21)), and secondary antibodies conjugated to Alexa 488, Cy3, or Cy5/AF647 (Jackson ImmunoResearch or Life Technologies; 1:400). All images of fixed samples were acquired using a DeltaVision Elite system (GE) equipped with a 100× 1.45 NA oil-immersion objective (Olympus) or a Marianas/SoRa spinning-disc confocal microscope (Intelligent Imaging Innovations, Inc.) at ambient temperature (21°C), using a 100× 1.46 NA oil immersion objective. Images were deconvolved using the SoftWoRx Suite (Applied Precision, GE) or the proprietary Microvolution software in Slidebook (Intelligent Imaging Innovations, Inc.) respectively. Identical imaging parameters (e.g., illumination power and exposure time) were used for all samples in the same experiment. All fixed gonads were imaged as 3D image stacks at intervals of 0.2 µm. Maximum-intensity projections of deconvolved 3D images were generated and tiled using Fiji unless otherwise noted. Figures were assembled using Adobe Illustrator.

### Live imaging

Live imaging was performed as previously described (45). Immobilized worms mounted on agarose pads were imaged with a Marianas/SoRa spinning-disc confocal microscope (Intelligent Imaging Innovations, Inc.) at ambient temperature (21°C), using a 100× 1.46 NA oil immersion objective. For apoptosis assays, differential interference contrast (DIC) and 488nm laser excitation were used to image and score AO-positive apoptotic nuclei in the germline. 3D images of the germline were acquired as Z-stacks of 26 focal planes at intervals of 0.5 µm, which spanned approximately half of the total thickness of tissue. For dynamics of the NE marker SUN-1::mRuby, 3D image stacks were acquired every 5 sec for a total duration of 5 minutes, with 12 Z-sections at intervals of 0.5 µm per time point. 488 or 561 nm excitation lasers were used with identical parameters between control and experimental samples. Deconvolved 3D images were assembled using Fiji and Illustrator (Adobe) for figures, unless otherwise noted.

### Image Analysis

Fluorescence intensity of FLAG (PLK-2) at HIM-8 foci or fluorescence intensity of LMN-1(V5) of each nucleus was manually segmented and measured from additive projection images after background subtraction using Fiji. For endogenous LMN-1 staining, due to limited signal-to-noise ratio, fluorescence intensity was measured along the circumference of the cross section of each nucleus with background subtraction in Fiji. Percentages of nuclei positive for LMN-1-pSer32 staining or types of distribution of fluorescently tagged PEZO-1 were measured manually. Quantification of the fraction of nuclei with complete synapsis was carried out as previously described (45). For curvature analysis, n = 11 (-Auxin) or 15 (+ Auxin) late pachytene nuclei (pooled from at least four animals in each condition) were analyzed for the deformation of NE contours, which was manually segmented for at least 20 consecutive frames acquired every 5 seconds using Fiji and subsequently processed in MATLAB (R2023a, Mathworks) (111). Curvature measurement at any given moment for each nucleus over the whole tracked duration are combined for plotting the histogram. Pixels that lie along NE contours were converted from Cartesian coordinates to polar coordinates. “Static corrugation” is the standard deviation over all polar angles of the contour radius for each nucleus at one time point. The histogram shows the distribution of this value for all time points. “Dynamic fluctuation” is the standard deviation of radii at each angular position for all time points in a series. The histogram shows the distribution of this value for all angles.

## Statistics

All statistical analyses were carried out using RStudio (Version 1.2.5033) or MATLAB (R2023a, Mathworks). All *P* values were calculated using the Mann-Whitney test or 2-sample t-test when analyzing the number of apoptotic nuclei per gonad arm using AO assays (Figures 1D, 1F, 1G; 2A, 2C to 2E; 5E; 6C; S3; S7; S9D). All other statistical test results are included in figure legends (except Figure 5B, which is included in “Image Analysis”).

## Acknowledgements

We thank members of the Dernburg Laboratory for stimulating discussions. We thank Zoe Lung and Xinyi Liu for their assistance. We thank Verena Jantsch (University of Vienna), Xiaofei Bai (University of Florida), Andy Golden (NIDDK), Needhi Bhalla (UC Santa Cruz), Yosef Gruenbaum (The Hebrew University of Jerusalem), and Yumi Kim (Johns Hopkins University) for reagents and advice. We also thank Anne Villeneuve (Stanford University), Ofer Rog (The University of Utah), Dan Starr (UC Davis), Regina Bohn (UC Davis), Amy Gladfelter and Dan Kiehart (Duke University), Kelly Liu and Mariana Wolfner (Cornell University), Rebecca Heald and Matt Welch (UC Berkeley) for helpful discussions. We are particularly grateful to Yu He (Yale University) for advice on quantification. Some *C. elegans* strains used in this work were provided by the Caenorhabditis Genetics Center, which is funded by the NIH – Office of Research Infrastructure Programs (P40 OD010440).

## Author contributions

Conceptualization: CL, AFD; Methodology: CL, AFD; Investigation: CL, AFD; Visualization: CL; Supervision: AFD; Project administration: AFD; Writing—original draft: CL; Writing—review editing: CL, AFD; Funding acquisition: CL, AFD.

## Data and materials availability

All data needed to evaluate the conclusions in the paper are present in the paper and/or the Supplementary Materials. Strains generated in this study can be provided upon request.

## Supplementary Materials

Figures S1 to S12; Tables S1 to S4; Movies S1 and S2.

**Figure S1.**
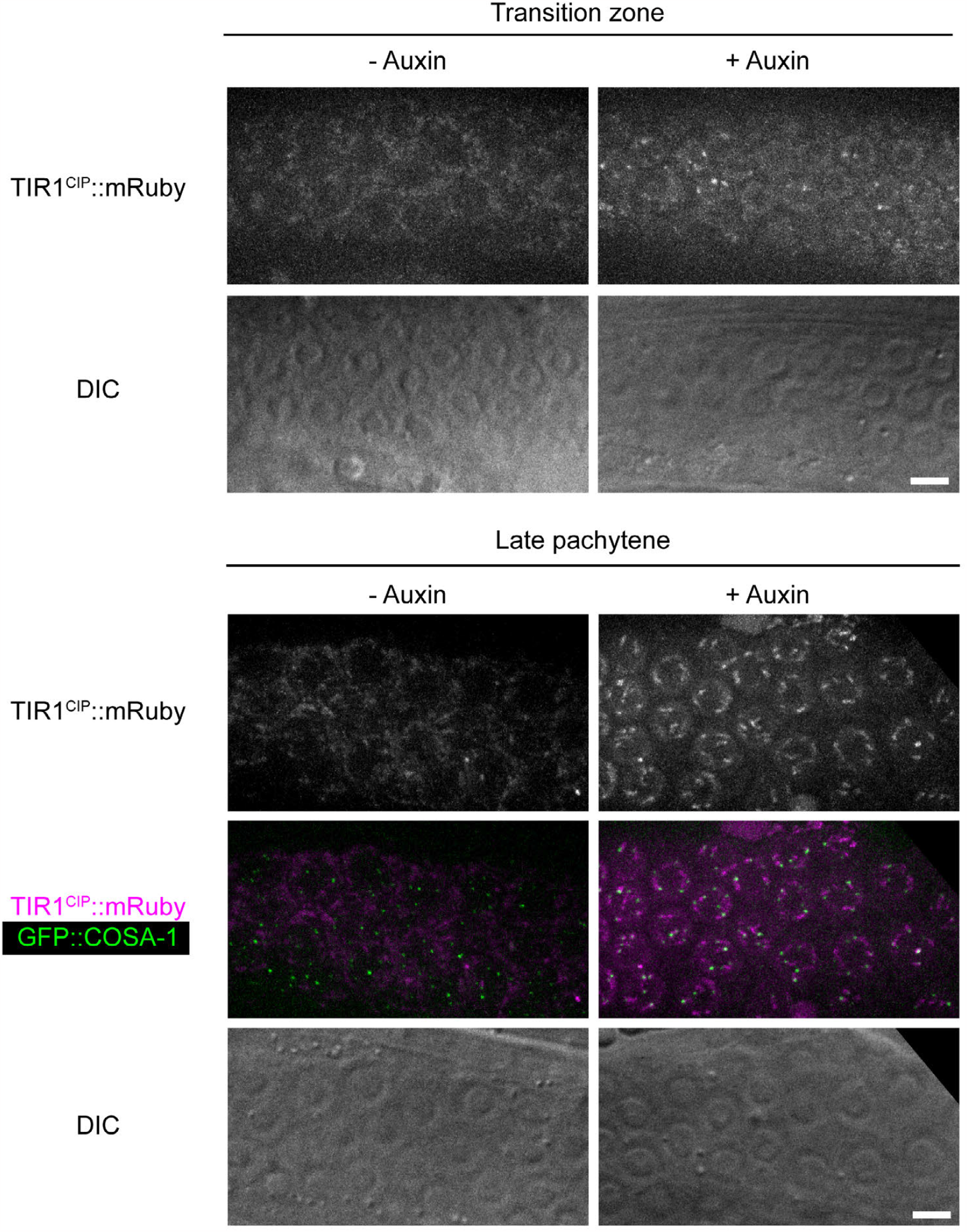
(Related to Figure 1) Repurposing the auxin-inducible degradation system into a chemically induced proximity system. Recruiting TIR1CIP::mRuby to degron-tagged PLK-2 results in fluorescence at pairing centers in transition zone nuclei and short arms of the SC at late pachytene, recapitulating prior localization of wild-type PLK-2. GFP::COSA-1 marks designated crossover sites that form boundaries between the long and short arms of the SC (1–3). Maximum-intensity projections of spinning-disk confocal data stacks are shown. Scale bars, 5 µm.

**Figure S2.**
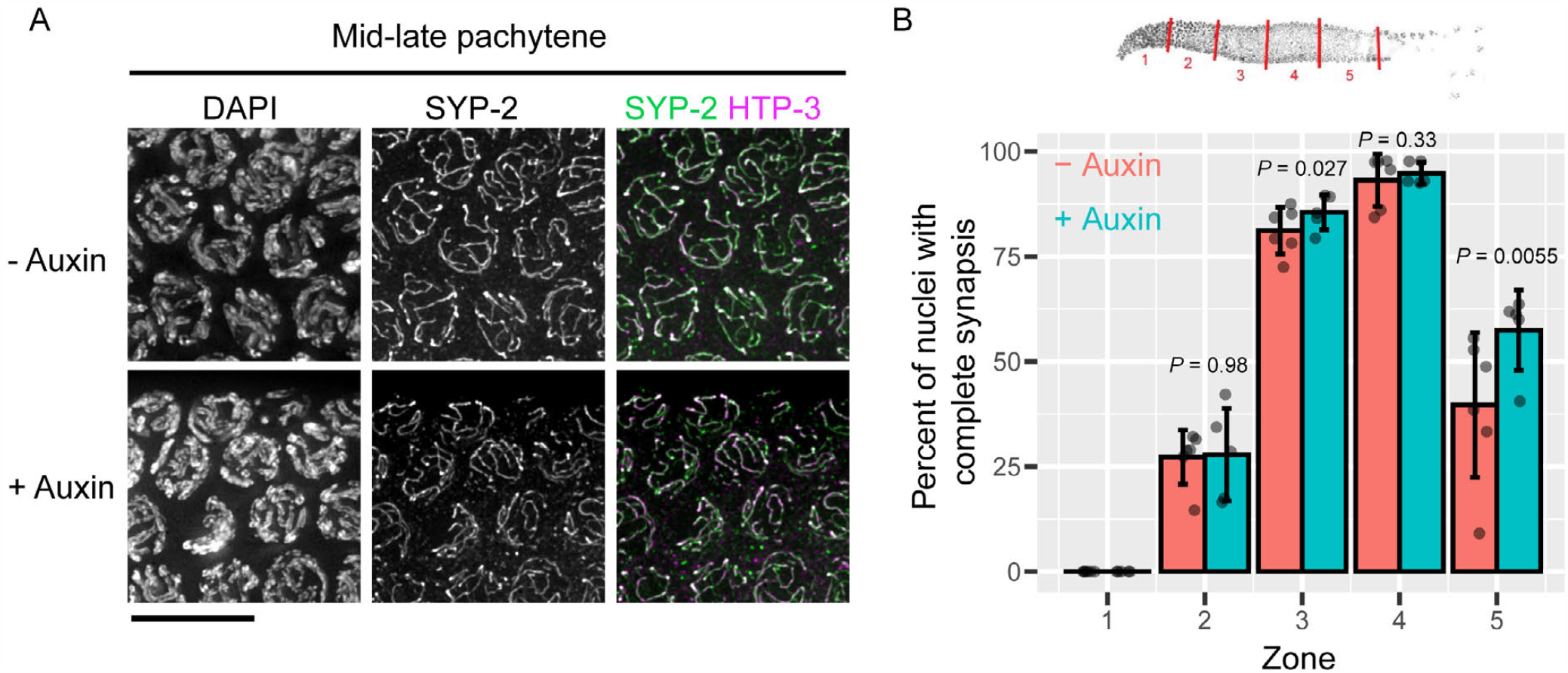
(Related to Figure 1) Recruiting PLK-2 to HIM-8 uncouples PLK-2 localization at pairing centers from asynapsis. (A) Representative immunofluorescence images showing mid to late-pachytene nuclei stained for SYP-2 (green) as an SC marker and HTP-3 (magenta) as an axis marker. Scale bar, 10 µm. (B) Quantification of nuclei with complete synapsis during meiotic progression. Diagram of distal gonad divided into five zones of equal length. Each dot represents one animal. Mean ± SD are plotted. Two-proportions z-test was used to compute the *P* values.

**Figure S3.**
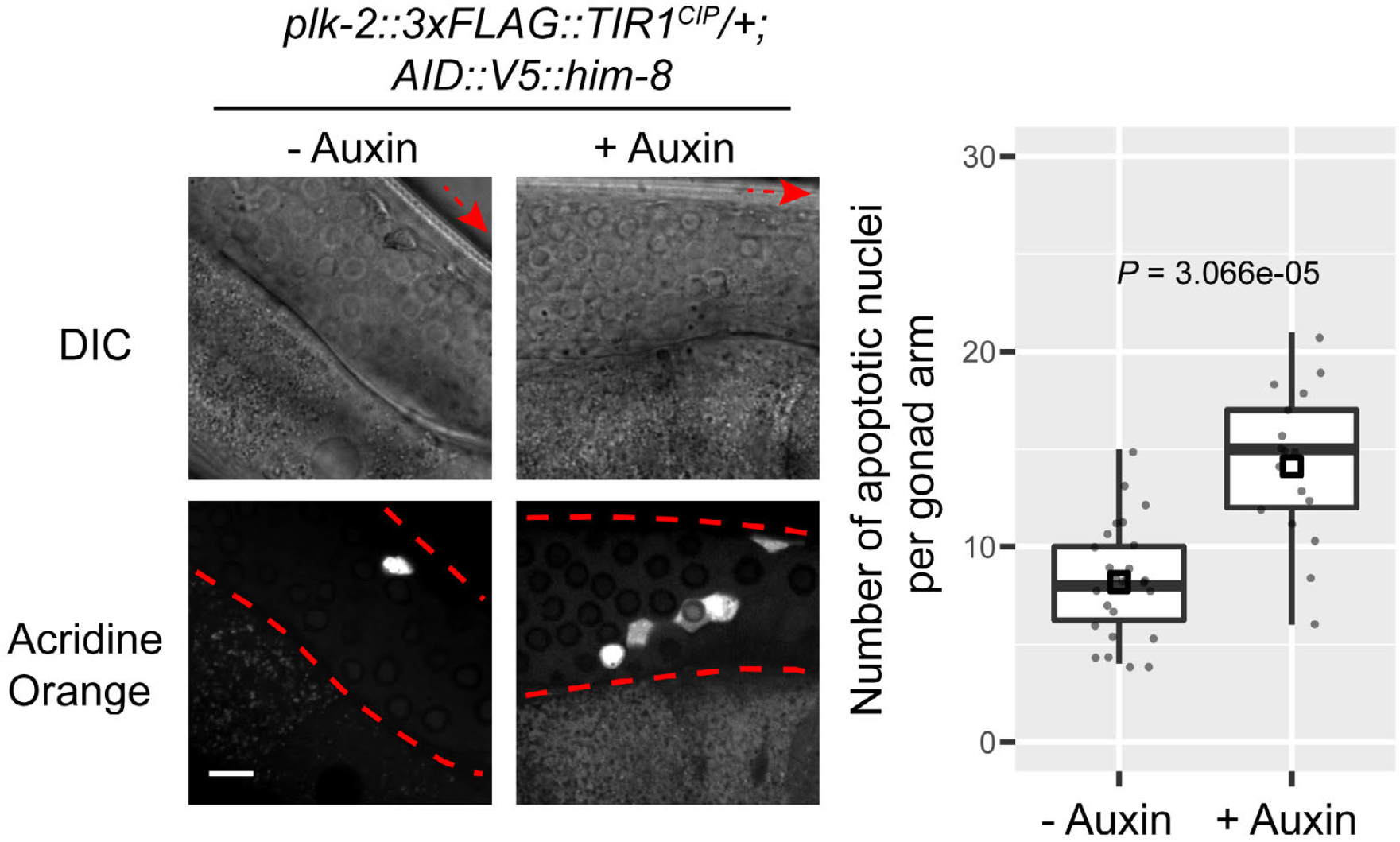
(Related to Figure 1) Targeting PLK-2 to X chromosome PCs triggers apoptosis even in the presence of untagged PLK-2. Representative images showing DIC and acridine orange staining of heterozygous animals carrying one copy of *plk-2::TIR1CIP* and one copy of wild-type *plk-2*, without or with auxin treatment. Scale bar, 10 µm. Boxplots show quantification of apoptotic nuclei based on AO staining. Each dot represents one animal. Medians (black crossbars) and means (black boxes) are shown.

**Figure S4.**
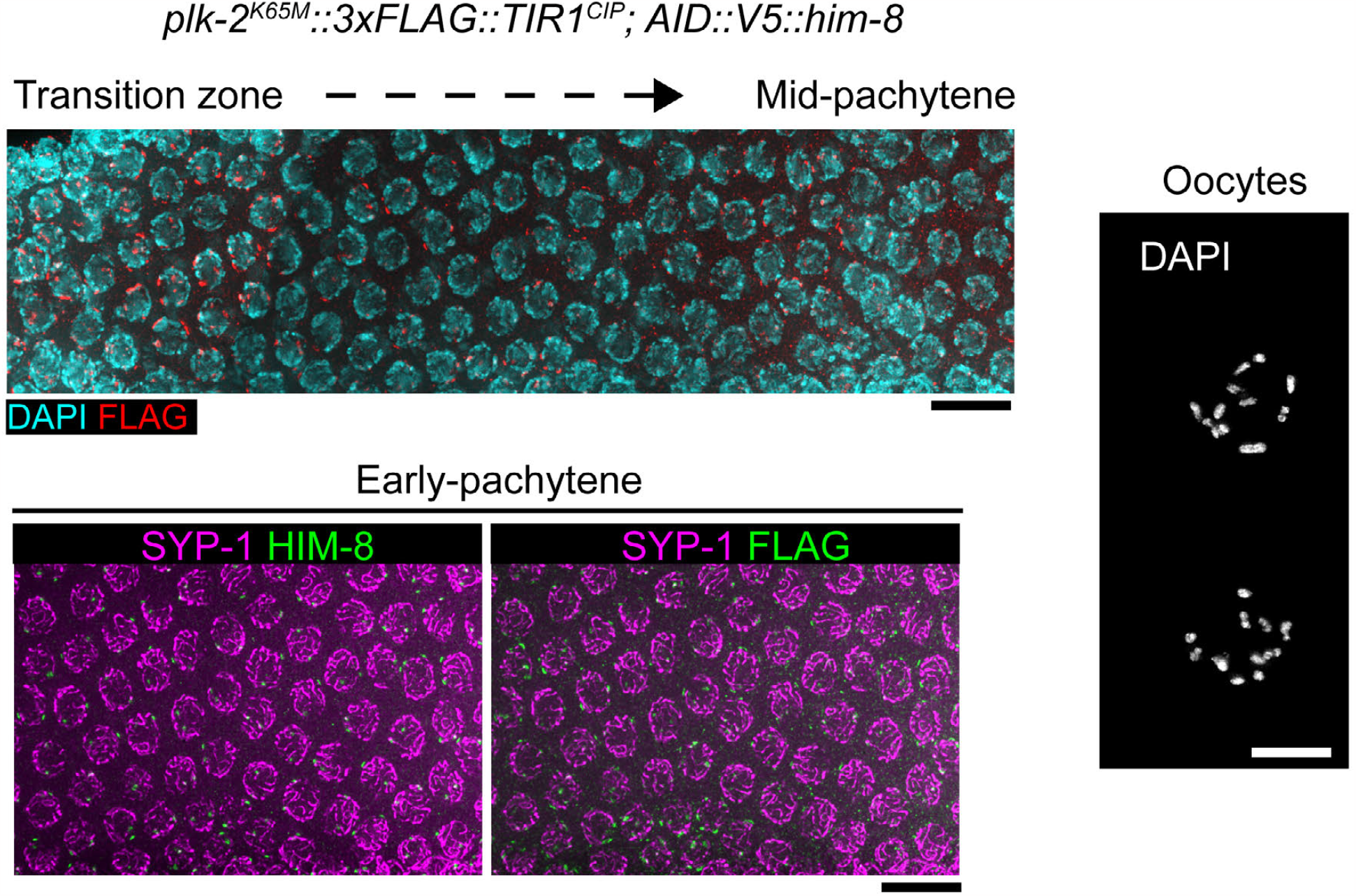
(Related to Figure 2) The kinase activity of PLK-2 is essential for meiosis. Representative immunofluorescence images showing phenotypes in meiotic nuclei from animals homozygous for *plk-2(K65M)::3xFLAG::TIR1CIP*. The absence of polarized nuclear morphology and defects in chromosome pairing and synapsis recapitulate phenotypes previously described in worms homozygous for untagged *plk-2(K65M)* (4; 5) or a similar *plk-2(P197L)* mutant defective in kinase activity (6). Scale bars, 10 µm.

**Figure S5.**
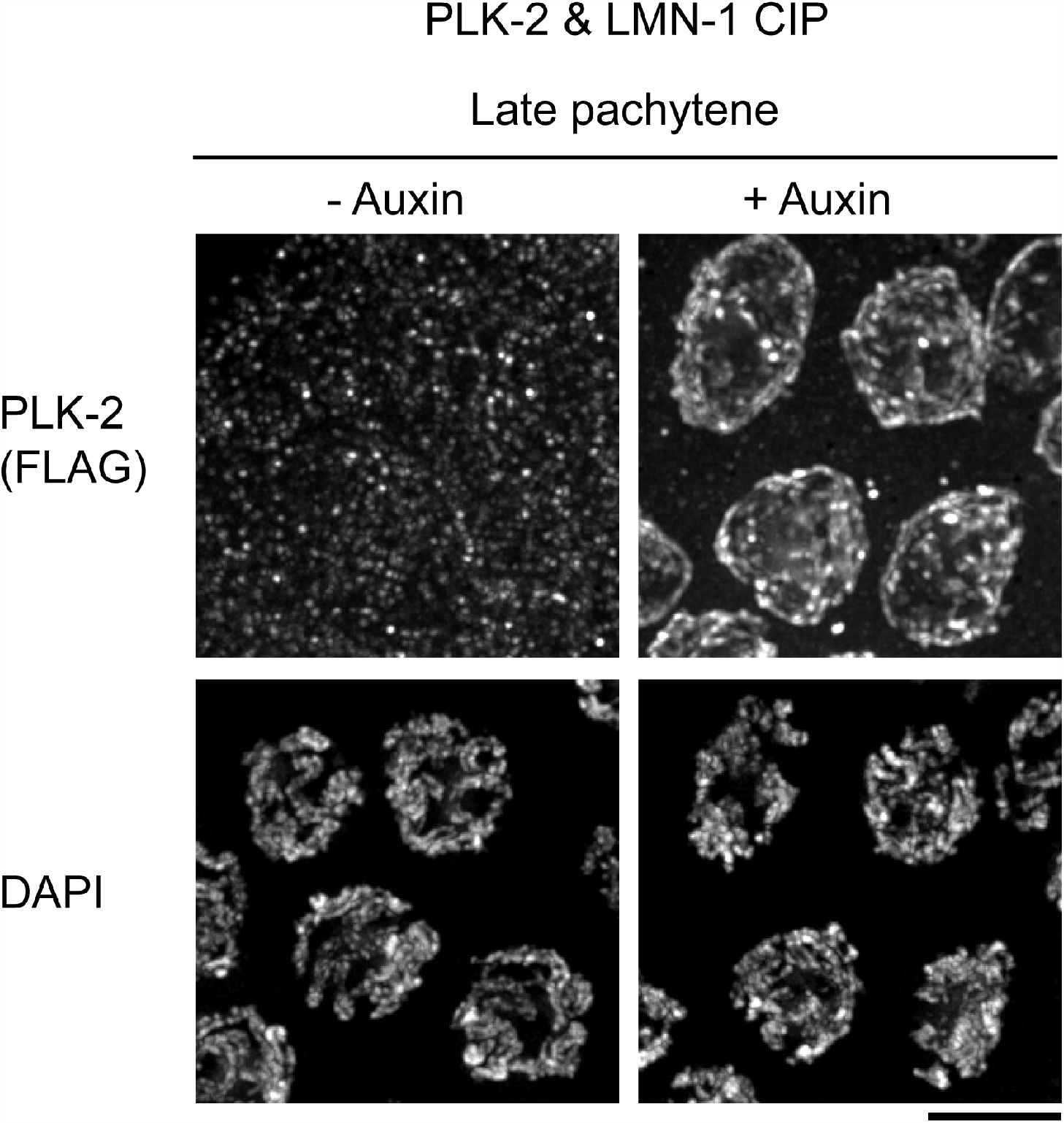
(Related to Figure 2) Recruiting PLK-2 to LMN-1 using CIP. Representative immunofluorescence showing late pachytene nuclei stained for DAPI and PLK-2 (FLAG), without or with auxin treatment to recruit PLK-2::3xFlAG::TIR1CIP to degron tagged LMN-1::V5. Scale bar, 5 µm.

**Figure S6.**
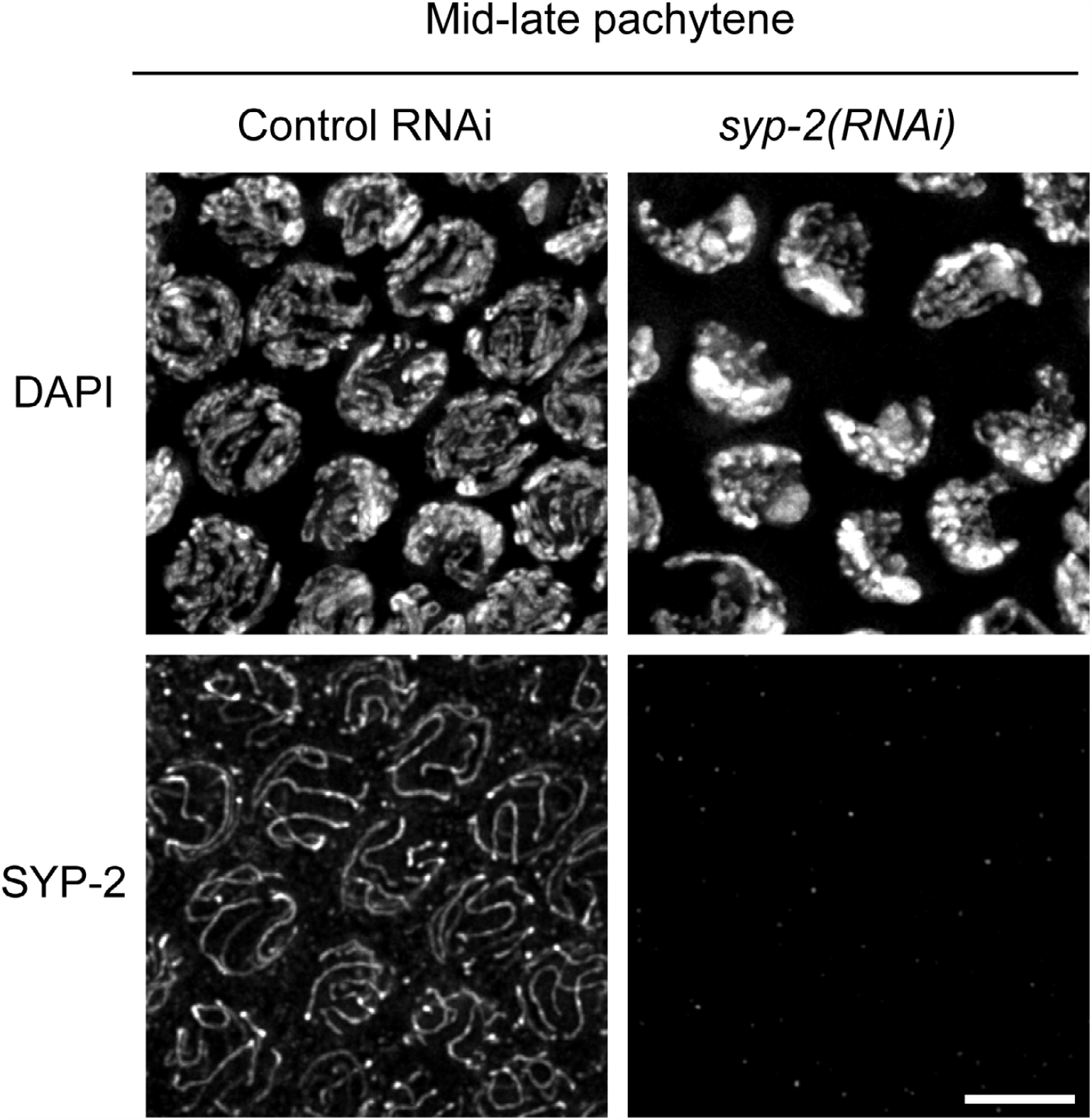
(Related to Figure 4) Depletion of SYP-2 by RNAi causes asynapsis. Representative immunofluorescence images showing mid to late pachytene nuclei stained for SYP-2 as an SC marker. DAPI counterstaining shows chromosome morphology. Scale bar, 5 µm.

**Figure S7.**
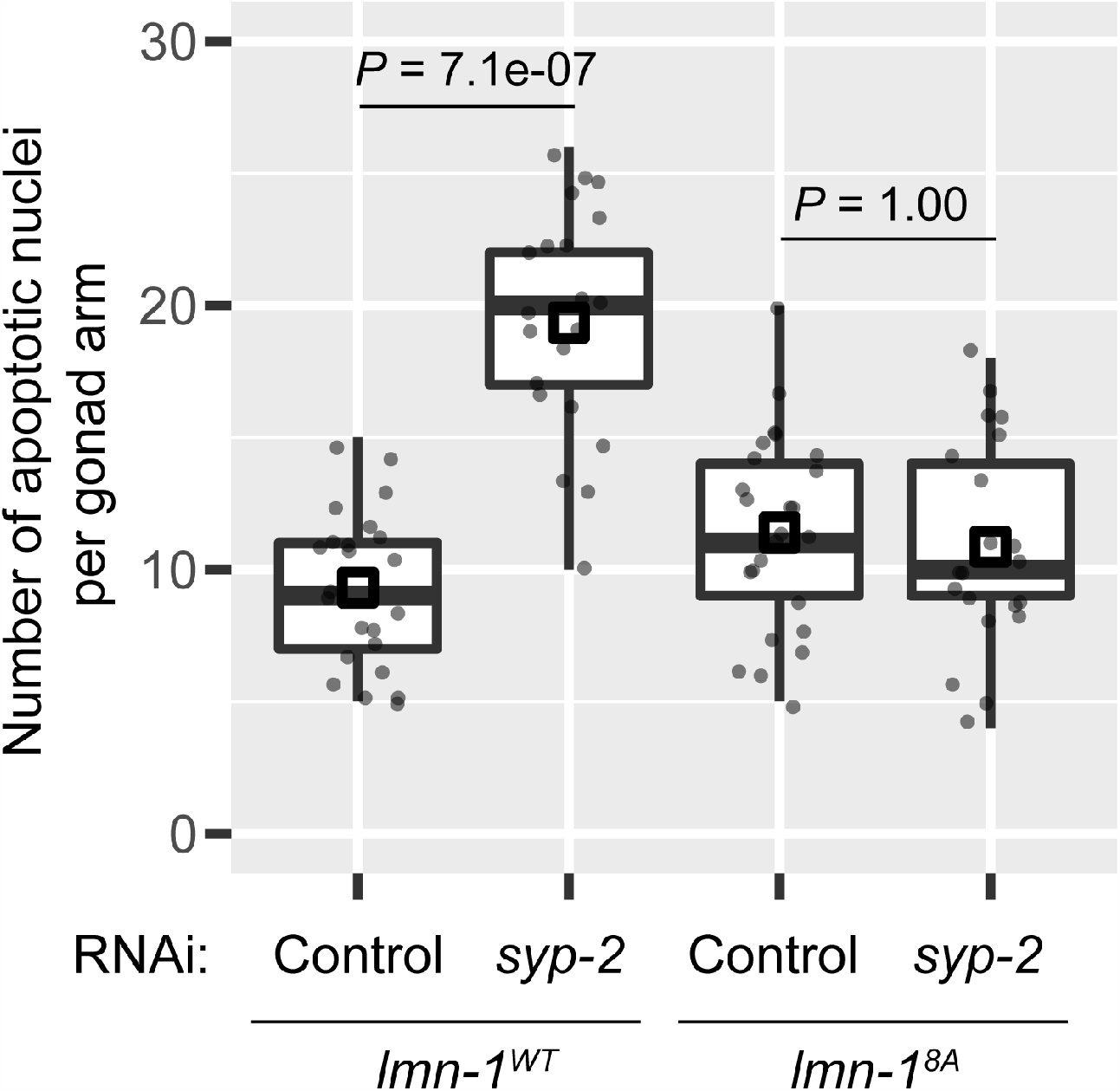
(Related to Figure 2 and Figure 4) *lmn-1*^*8A*^ abolishes elevated apoptosis in response to asynapsis. Unlike SYP-2 depletion in the wild-type *lmn-1* background, SYP-2 depletion in the nonphosphorylatable *lmn-1*^*8A*^ background fails to trigger elevated apoptosis. Boxplots show quantification of apoptotic nuclei based on AO staining. Each dot represents one animal. Medians (black crossbars) and means (black boxes) are shown.

**Figure S8.**
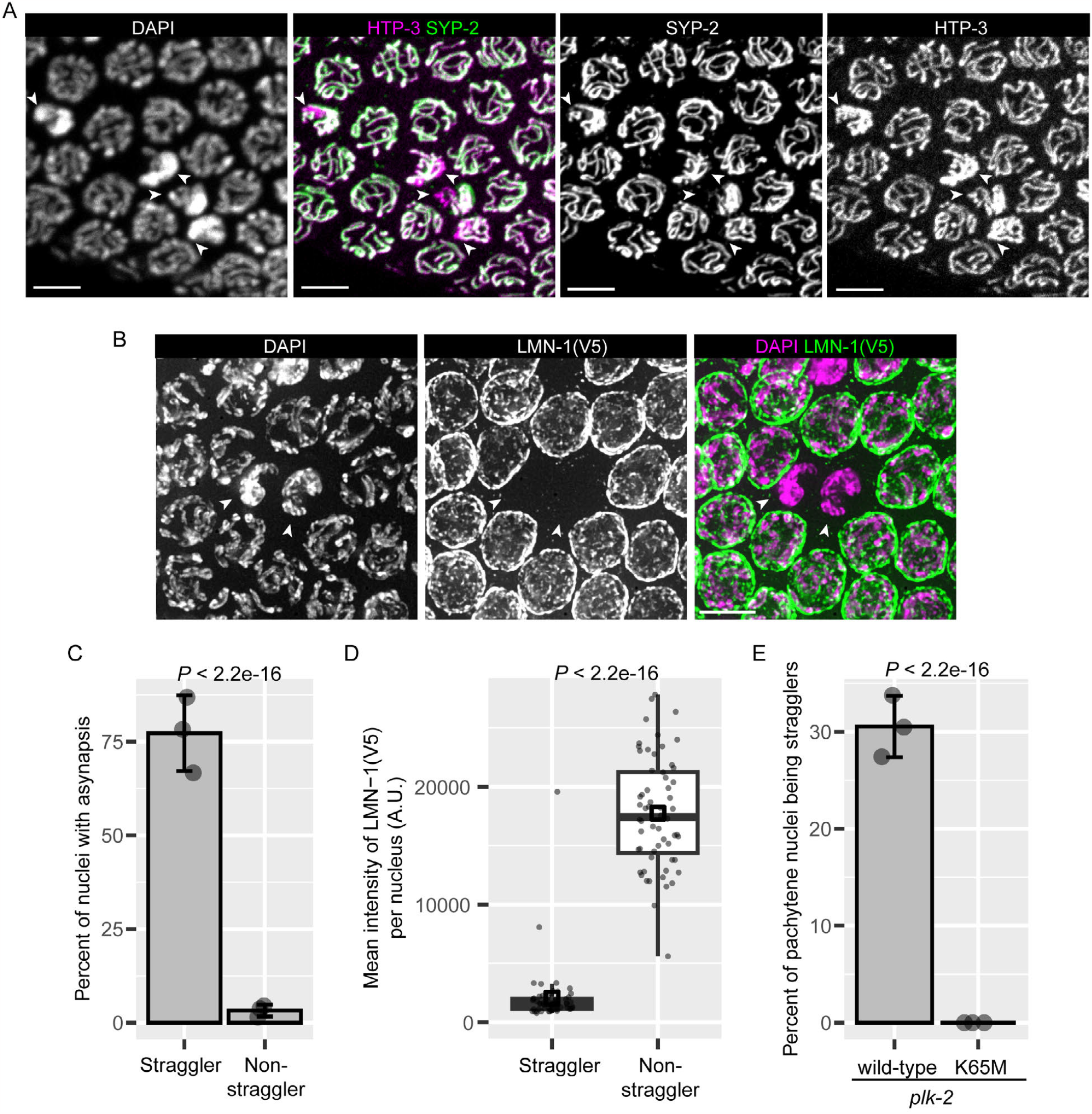
(Related to Figure 3 and Figure 4) “Straggler” nuclei detected in wild-type animals have weakened nuclear lamina. (A) Representative immunofluorescence images showing “straggler” nuclei in mid-pachytene with a characteristic clustered and polarized chromosomal arrangement and defective synapsis. SC is marked by SYP-2 and chromosomal axes by HTP-3. Arrowheads point to “straggler” nuclei. Maximum-intensity projections of raw images acquired on a spinning-disk confocal are shown. Scale bars, 5 µm. (B) Representative immunofluorescence images showing “straggler” nuclei having weakened nuclear lamina, indicated by arrowheads. Scale bar, 5 µm. (C) “Stragglers” often display incomplete synapsis. Bar plot shows the percentage of nuclei with asynapsis (segments of chromosomes positive for HTP-3 but negative for SYP-2) among “stragglers” and “non-stragglers”. Mean ± SD from three animals is plotted. *P* was computed using the two-proportions z-test. The population of meiotic nuclei analyzed was in mid-pachytene, prior to desynapsis at diplotene. (D) Staining of the nuclear lamina in straggler nuclei is much fainter than in adjacent pachytene nuclei. Boxplots show quantification of mean intensity of LMN-1::V5 per nucleus. 20 pachytene nuclei from each of 3 animals were measured per condition. Medians (black crossbars) and means (black boxes) are shown. *P* value was computed using two sample t-test. (E) PLK-2 kinase activity is required for the appearance of stragglers. Bar plot shows the percentage of straggler nuclei in all mid-pachytene nuclei. Mean ± SD from three animals is plotted. *P* was computed using the two-proportions z-test.

**Figure S9.**
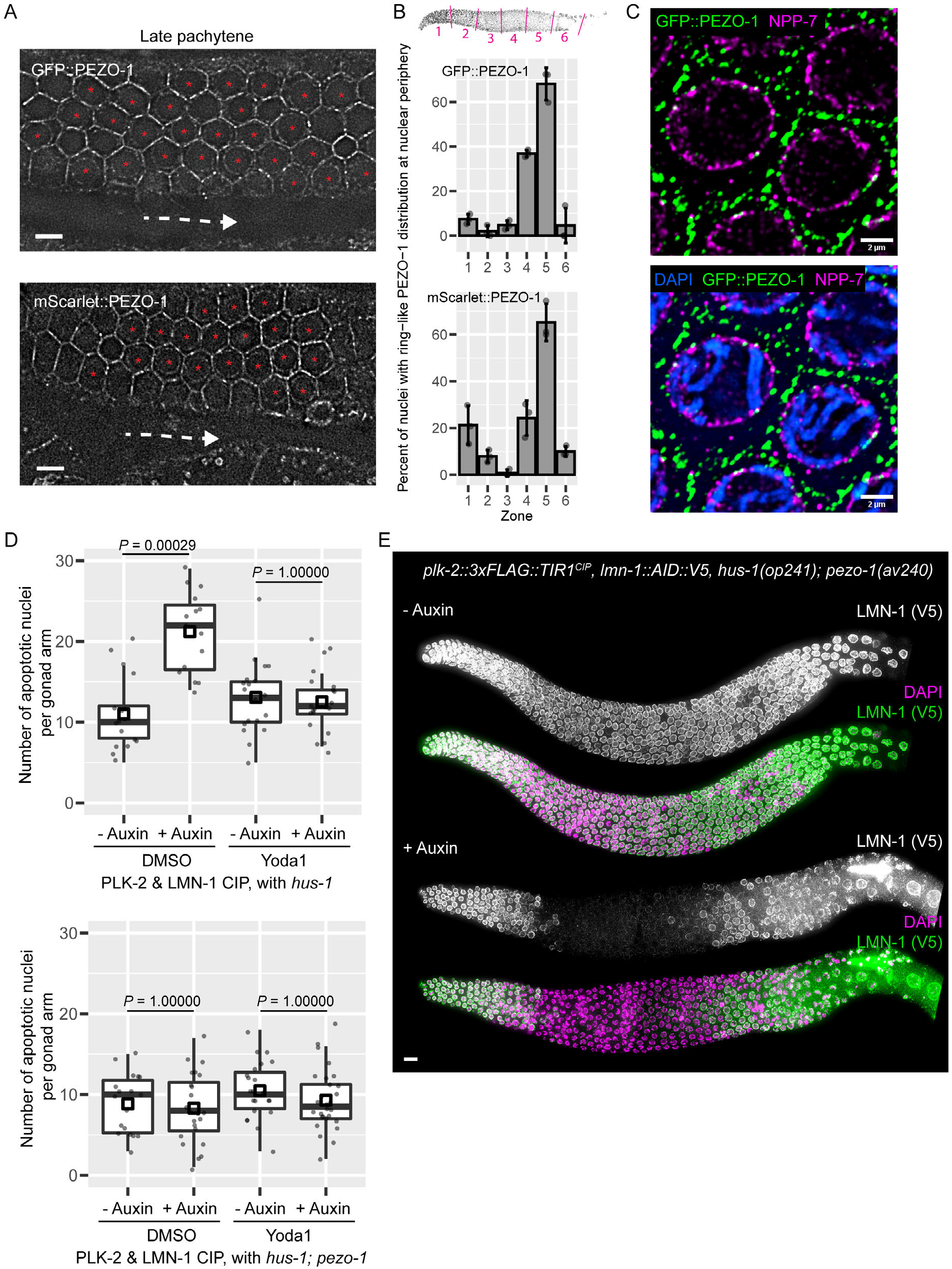
(Related to Figure 5) PEZO-1 localizes to the nuclear periphery in late pachytene and is important for the synapsis checkpoint. (A) Confocal images of gonads in live hermaphrodites expressing GFP::PEZO-1 or mScarlet::PEZO-1. Dashed white arrow indicates the direction of meiotic progression. Red asterisks indicate meiotic cells with “ring-like” localization of PEZO-1 at the nuclear periphery. Scale bars, 5 µm. (B) Quantification of nuclei with “ring-like” PEZO-1 distributions at the nuclear periphery. Diagram of distal gonad divided into six zones of equal length. Gonads from three animals were measured per condition. Mean ± SD are plotted. (C) Representative immunofluorescence images showing GFP::PEZO-1 and NPP-7 (Nup153 (7)) at the NE in late pachytene. Fixed samples were imaged using a SoRa spinning disk confocal. Single optical sections from deconvolved 3D data stacks are shown. Scale bars, 2µm. (D) Elevated apoptosis following recruitment of PLK-2 to the nuclear lamina is abrogated by a Piezo1 agonist. Boxplots show quantification of apoptotic nuclei based on AO staining. Each dot represents one animal. Medians (black crossbars) and means (black boxes) are shown. (E) Representative images showing a lamin “dark zone” following PLK-2 recruitment to the nuclear lamina in animals lacking PEZO-1. Scale bar, 10 µm.

**Figure S10.**
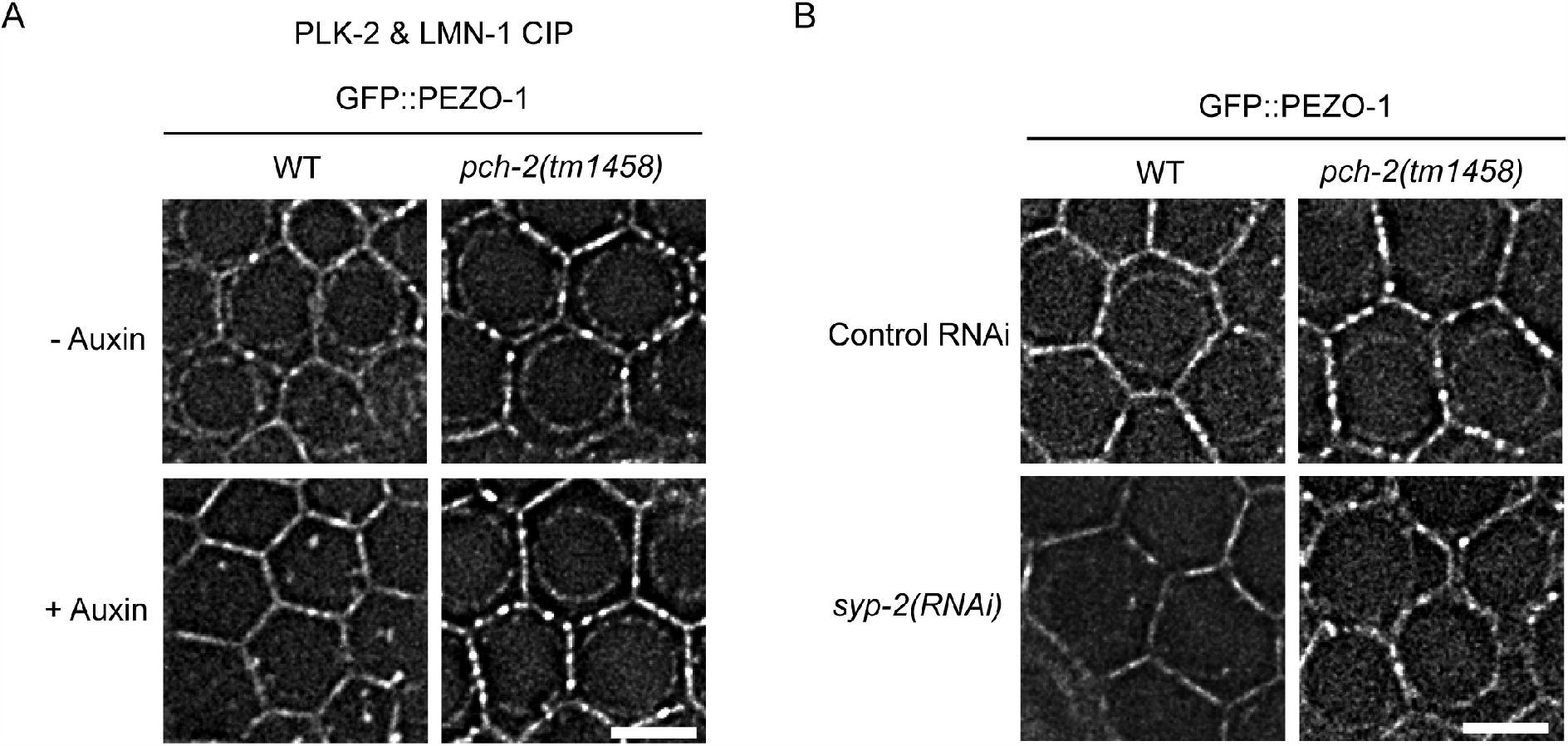
(Related to Figure 6) PEZO-1 redistributes at the NE in a PCH-2 dependent manner. Representative frames showing gonads in live worms expressing GFP::PEZO-1, which is seen at the plasma membrane and nuclear periphery in both wild-type and *pch-2(tm1458)* hermaphodites. (A) GFP::PEZO-1 localization in gonad following the recruitment of PLK-2 to LMN-1. (B) GFP::PEZO-1 localization in gonad following control or *syp-2(RNAi)*. Scale bars, 5 µm.

**Figure S11.**
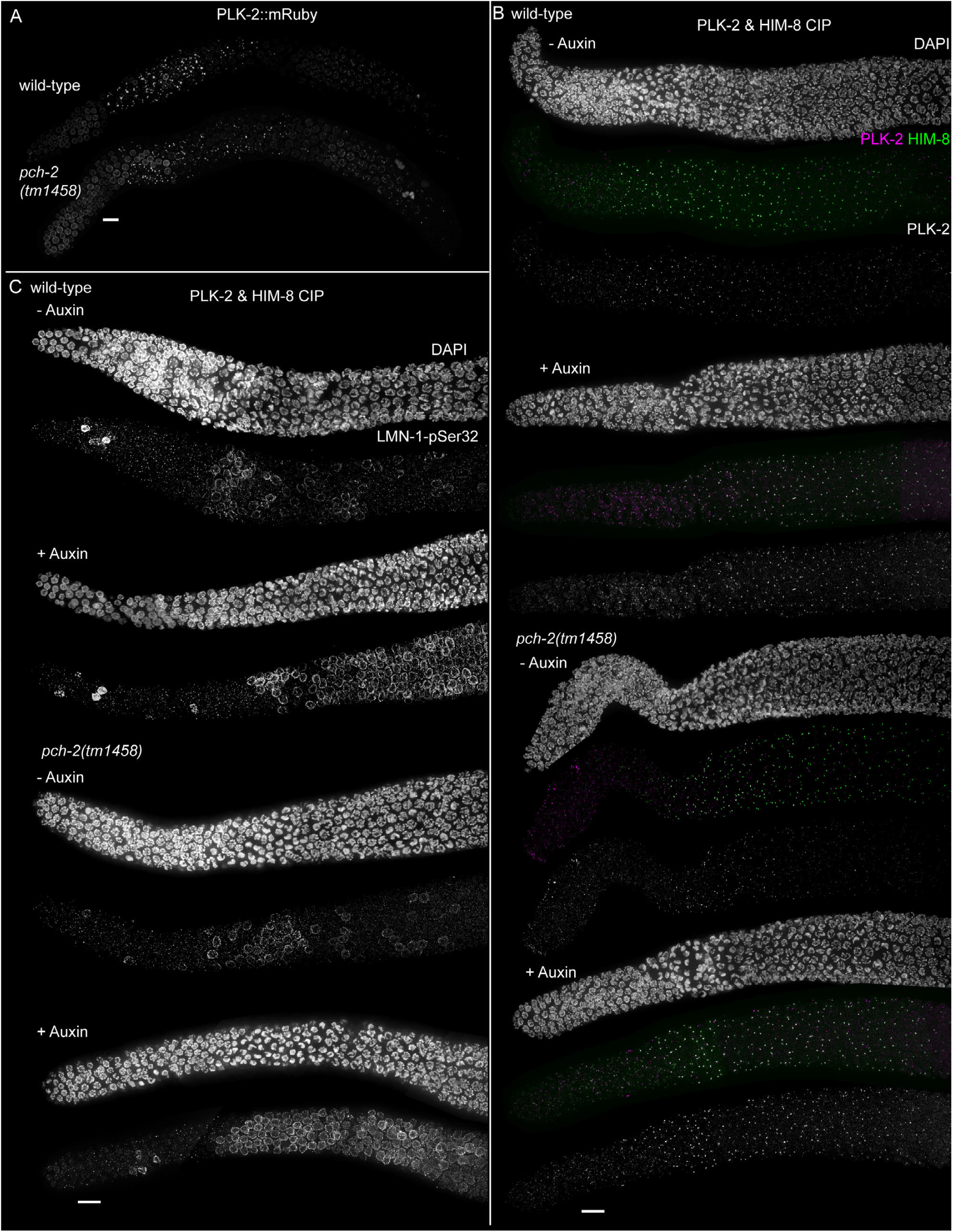
(Related to Figure 6) PCH-2 is not required for the localization or activity of PLK-2 in response to asynapsis. (A) PLK-2::mRuby still localizes to PCs at the NE of transition zone nuclei in *pch-2(tm1458)* animals. Maximum-intensity projections of 3D data stacks from live animals are shown. (B) Upon auxin treatment to recruit PLK-2 to PCs, the extended localization of PLK-2::3xFlAG::TIR1CIP at PCs appears indistinguishable in both wild-type and *pch-2(tm1458)* animals. (C) Upon auxin treatment to recruit PLK-2 to PCs, extended staining of LMN-1-pSer32 is observed in both wild-type and *pch-2(tm1458)* animals. (B) and (C) show representative immunofluorescence images. Scale bars, 10 µm.

**Figure S12.**
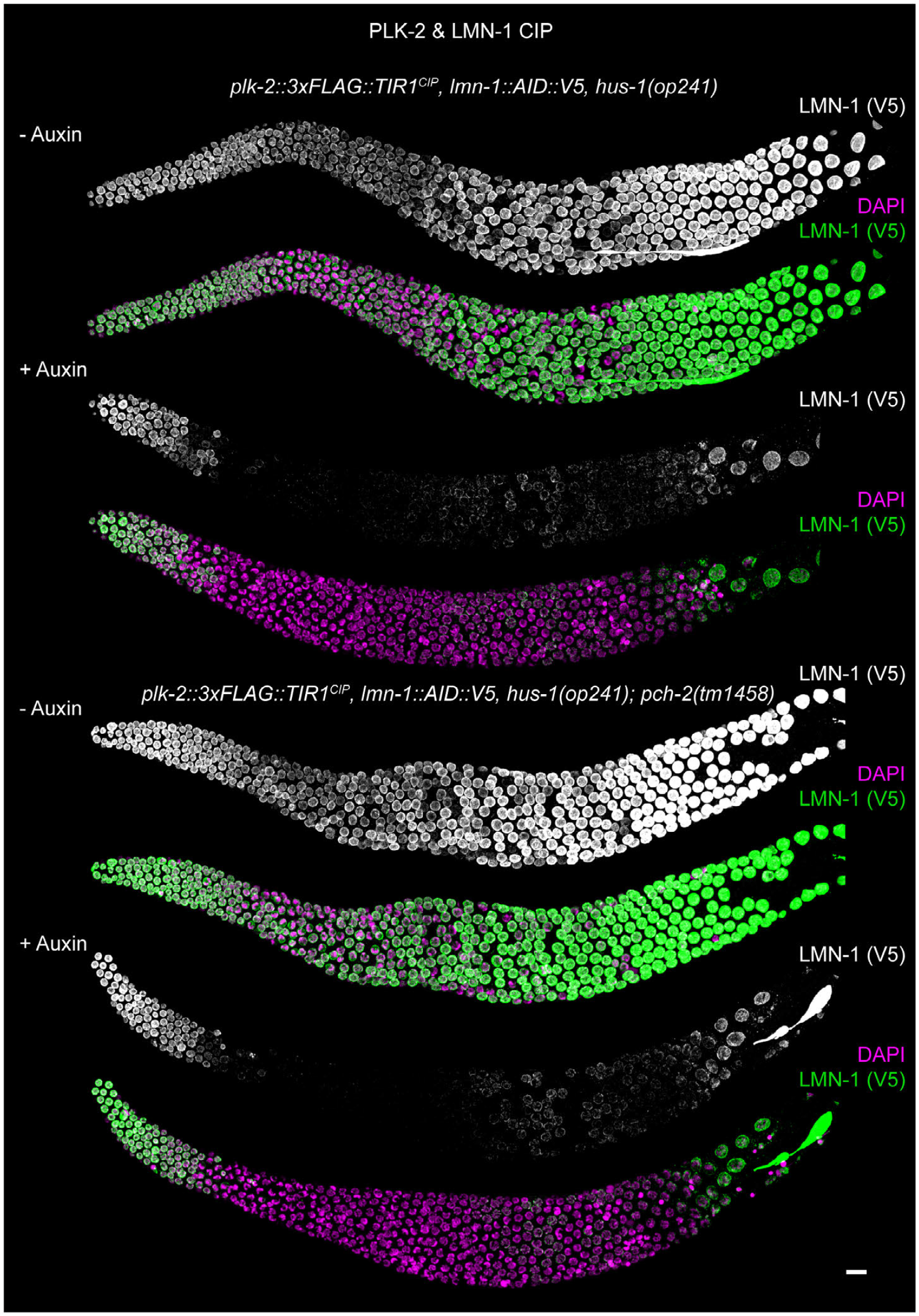
(Related to Figure 6) PCH-2 is not required for the lamin “dark zone” caused by PLK-2 recruitment to the nuclear lamina. Representative immunofluorescence images from *hus-1(op241)* and *hus-1(op241)*; *pch-2(tm1458)* animals following recruitment of PLK-2 to LMN-1. Scale bar, 10 µm.

## Supplementary Movies

**Movie S1.**
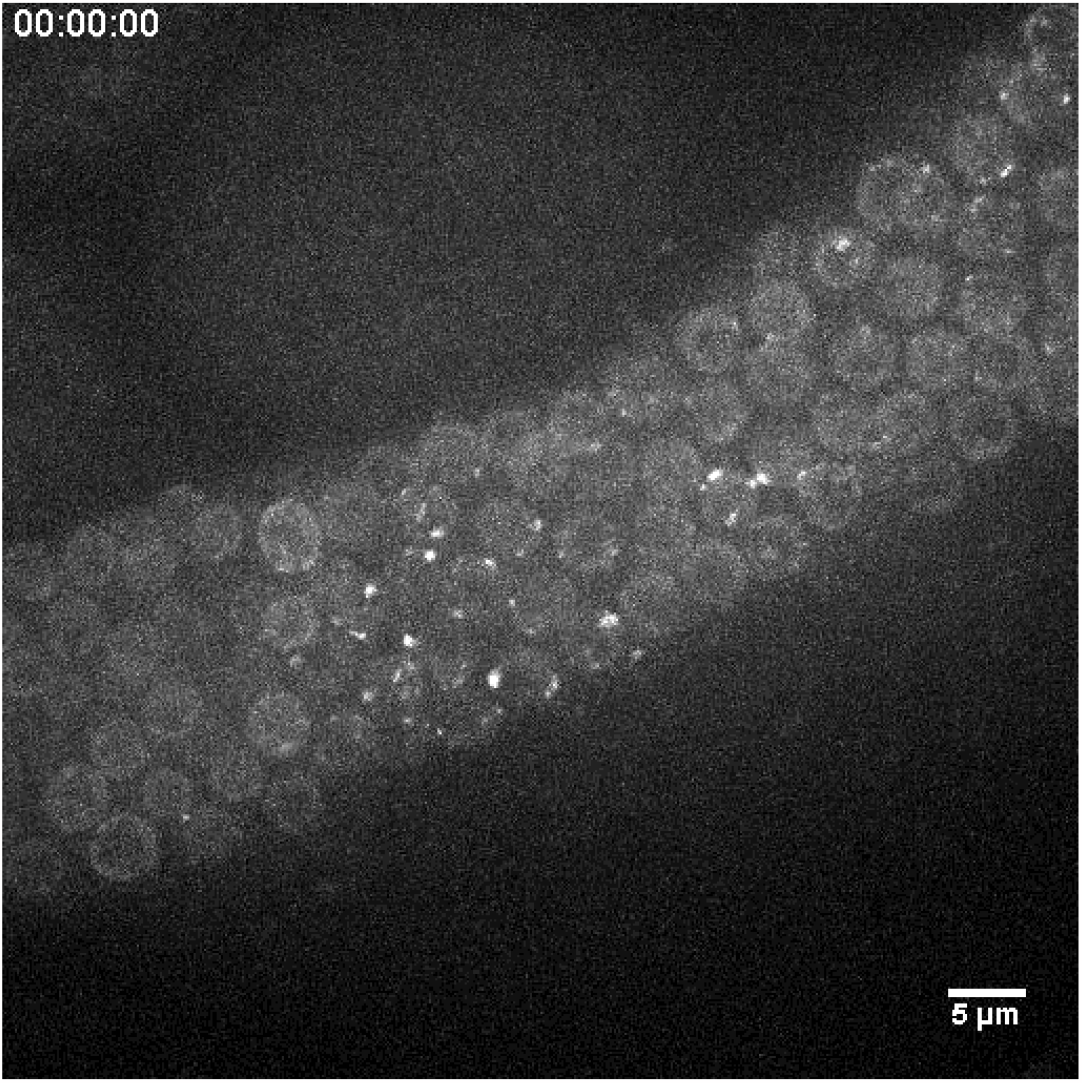
Recruiting TIR1CIP::mRuby to degron-tagged PLK-2 reveals PLK-2 dynamics at the NE. Time-lapse recording of a representative gonad from an intact live animal shows saltatory movements of pairing centers at the NE of transition zone nuclei. Meiosis proceeds from left to right. Time stamp is hr:min:sec. Scale bar, 5 µm.

**Movie S2.**
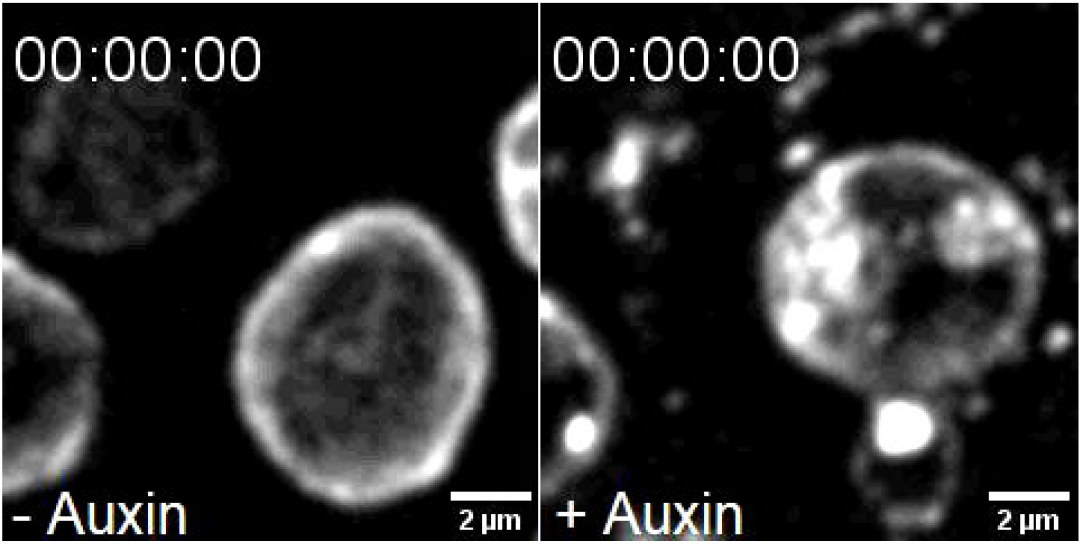
Recruiting PLK-2 to the nuclear lamina destabilizes the NE. The NE is visualized by expression of SUN-1::mRuby. Side-by-side comparison of time-lapse recordings of NE fluctuations in late pachytene nuclei, with or without auxin treatment. Time stamp is hr:min:sec. Scale bars, 2 µm.

**Table S1.** Alleles generated in this study.

| **Allele** | **Genotype** | **Information about mutagenesis** |
| --- | --- | --- |
| *ie204* | *TIR1<sup>CIP</sup>::mRuby* (transgene) | generated using *dpy-10* Co-CRISPR in *plk-2(ie108[plk-2::AID::3xFLAG])*, *zhp-2(ie107[zhp-2::HA]) I*; *mels8[pie-1p::GFP::cosa-1, unc-119(+)] II*; *ieSi38[sun-1p::TIR1::mRuby::sun-1 3'UTR, Cbr-unc-119(+)] IV* |
| *ie205* | *plk-2(ie205[plk-2::3xFLAG::TIR1<sup>CIP</sup>])* | generated using *dpy-10* Co-CRISPR in N2 wild-type |
| *ie206* | *plk-2(ie206[plk-2(K65M):3xFLAG::TIR1<sup>CIP</sup>])* | generated using *dpy-10* Co-CRISPR in *plk-2(ie205[plk-2::3xFLAG::TIR1<sup>CIP</sup>]) I*, *him-8(ie207[AID::V5::him-8]) IV* |
| *ie207* | *him-8(ie207[AID::V5::him-8])* | generated using *dpy-10* Co-CRISPR in N2 wild-type |

**Table S2.** Genotypes of worm strains generated and used in this study.

| **Strain** | **Source** | **Identifier** |
| --- | --- | --- |
| *C. elegans*: N2 Bristol, wild isolate | Caenorhabditis Genetics Center | N2 |
| *C. elegans*: *plk-2*(*ie108*[*plk-2*::*AID*::3xFLAG]), *zhp-2*(*ie107*[*zhp-2*::HA]) I; *mels8*[*pie-1p*::GFP::*cosa-1*, *unc-119*(+)] II; *ieSi38*[*sun-1p*::*TIR1*::*mRuby*::*sun-1* 3'UTR, *Cbr-unc-119*(+)] IV | Zhang et al. (1);<br>Caenorhabditis Genetics Center | CA1429 |
| *C. elegans*: *plk-2*(*ie108*[*plk-2*::*AID*::3xFLAG]), *zhp-2*(*ie107*[*zhp-2*::HA]) I; *mels8*[*pie-1p*::GFP::*cosa-1*, *unc-119*(+)] II; *ieSi38*[*sun-1p*::*TIR1*<sup>CIP</sup>::*mRuby*::*sun-1* 3'UTR, *Cbr-unc-119*(+)] IV | This paper | CA1767 |
| *C. elegans*: *plk-2*(*ie205*[*plk-2*::3xFLAG::*TIR1*<sup>CIP</sup>]) I; *him-8*(*ie207*[*AID*::V5::*him-8*]) IV | This paper | CA1768 |
| *C. elegans*: *hus-1*(*op241*) I | Bhalla et al. (2);<br>Caenorhabditis Genetics Center | CA125<br>(WS2277) |
| *C. elegans*: *pch-2*(*tm1458*) II | Bhalla et al. (2);<br>Japanese National Bioresource for *C. elegans* | CA388 |
| *C. elegans*: *plk-2*(*ie205*[*plk-2*::3xFLAG::*TIR1*<sup>CIP</sup>]), *hus-1*(*op241*) I; *him-8*(*ie207*[*AID*::V5::*him-8*]) IV | This paper | CA1769 |
| *C. elegans*: *plk-2*(*ie205*[*plk-2*::3xFLAG::*TIR1*<sup>CIP</sup>]) I; *pch-2*(*tm1458*) II; *him-8*(*ie207*[*AID*::V5::*him-8*]) IV | This paper | CA1770 |
| *C. elegans*: *tmC20* [*unc-14*(*tmls1219*) *dpy-5*(*tm9715*)] I | Dejima et al. (3);<br>Caenorhabditis Genetics Center | FX30179 |
| *C. elegans*: *plk-2*(*ie206*[*plk-2*(K65M)::3xFLAG::*TIR1*<sup>CIP</sup>])/ *tmC20* I; *him-8*(*ie207*[*AID*::V5::*him-8*]) IV | This paper | CA1771 |
| *C. elegans*: *lmn-1*(*jf140*[S21,22,24,32,397,398, 403,405A]) I/hT2 [*bli-4*(*e937*) *let-?*(*q782*) *qls48*] (I;III) | Velez-Aguilera et al. (4) | UV2059 |
| *C. elegans*: *plk-2*(*ie205*[*plk-2*::3xFLAG::*TIR1*<sup>CIP</sup>]), *lmn-1*(*jf140*[S21,22,24,32,397,398, 403,405A]) I/hT2 [*bli-4*(*e937*) *let-?*(*q782*) *qls48*] (I;III); *him-8*(*ie207*[*AID*::V5::*him-8*]) IV | This paper | CA1772 |
| *C. elegans*: *plk-2*(*ie205*[*plk-2*::3xFLAG::*TIR1*<sup>CIP</sup>]) I/hT2 [*bli-4*(*e937*) *let-?*(*q782*) *qls48*] (I,III); *him-8*(*ie207*[*AID*::V5::*him-8*]) IV | This paper | CA1773 |
| *C. elegans*: *lmn-1*(*ie137*[*lmn-1*::*AID*::V5]) I; *unc-119* (*ed3*) III; *ieSi38* [*Psun-1*::*TIR1*::*mRuby*::*sun-1* 3'UTR, *cb-unc-119*(+)] IV | Liu et al. (5) | CA1532 |
| *C. elegans: plk-2(ie205[plk-2::3xFLAG::TIR1<sup>CIP</sup>]), lmn-1(ie137[lmn-1::AID::V5]) I* | This paper | CA1774 |
| *C. elegans: plk-2(ie205[plk-2::3xFLAG::TIR1<sup>CIP</sup>]), lmn-1(ie137[lmn-1::AID::V5]), hus-1(op241) I* | This paper | CA1775 |
| *C. elegans: hus-1(op241) I; ieDf2/mls11 IV; bcls39 (ced-1::GFP) V* | Harper et al. (6) | CA971 |
| *C. elegans: plk-2(ie205[plk-2::3xFLAG::TIR1<sup>CIP</sup>]), lmn-1(ie137[lmn-1::AID::V5]), hus-1(op241) I; ieDf2/mls11 IV; bcls39 (ced-1::GFP) V* | This paper | CA1776 |
| *C. elegans: plk-2(ie206[plk-2(K65M)::3xFLAG::TIR1<sup>CIP</sup>]), lmn-1(ie137[lmn-1::AID::V5])/tmC20 I* | This paper | CA1777 |
| *C. elegans: plk-2(ie205[plk-2::3xFLAG::TIR1<sup>CIP</sup>]), lmn-1(ie137[lmn-1::AID::V5]) I; ced-4(n1162)III* | This paper | CA1778 |
| *C. elegans: plk-2(ie205[plk-2::3xFLAG::TIR1<sup>CIP</sup>]), lmn-1(ie137[lmn-1::AID::V5]), hus-1(op241) I; ieSi21 (sun-1::mRuby) IV* | This paper | CA1779 |
| *C. elegans: pezo-1(av142[mScarlet::pezo-1]) IV* | Bai et al. (7) | AG404 |
| *C. elegans: pezo-1(av146 [GFP::pezo-1]) IV* | Bai et al. (7) | AG408 |
| *C. elegans: dhc-1(he250[mCherry::dhc-1]) I* | Schmidt et al. (8) | SV1619 |
| *C. elegans: dhc-1(he250[mCherry::dhc-1]) I; pezo-1(av146 [GFP::pezo-1]) IV* | This paper | CA1780 |
| *C. elegans: lmn-1(jf98[GFP::lmn-1]) I* | Link et al. (9) | UV117 |
| *C. elegans: lmn-1(jf98[GFP::lmn-1]) I; pezo-1(av142[mScarlet::pezo-1]) IV* | This paper | CA1781 |
| *C. elegans: pezo-1(av240) IV* | Bai et al. (7) | AG570 |
| *C. elegans: plk-2(ie205[plk-2::3xFLAG::TIR1<sup>CIP</sup>]), lmn-1(ie137[lmn-1::AID::V5]), hus-1(op241) I; pezo-1(av240) IV* | This paper | CA1782 |
| *C. elegans: plk-2(ie205[plk-2::3xFLAG::TIR1<sup>CIP</sup>]), lmn-1(ie137[lmn-1::AID::V5]), hus-1(op241) I; pezo-1(av146 [GFP::pezo-1]) IV* | This paper | CA1783 |
| *C. elegans: plk-2(ie205[plk-2::3xFLAG::TIR1<sup>CIP</sup>]), lmn-1(ie137[lmn-1::AID::V5]), hus-1(op241) I; pch-2(tm1458) II; pezo-1(av146 [GFP::pezo-1]) IV* | This paper | CA1784 |
| *C. elegans: dhc-1(he250[mCherry::dhc-1]) I; pch-2(tm1458) II; pezo-1(av146 [GFP::pezo-1]) IV* | This paper | CA1785 |
| *C. elegans: dhc-1(he264[GFP::dhc-1]) I* | Schmidt et al. (8) | SV1803 |
| *C. elegans: dhc-1(he264[GFP::dhc-1]) I; ieSi21 (sun-1::mRuby) IV* | This paper | CA1786 |
| *C. elegans: ieSi64[gld-1p::TIR1::mRuby::gld-1 3'UTR, Cbr-unc-119(+)] II; unc-119(ed3) III; sun-1(ie139[sun-1::AID::V5]) V* | This paper | CA1787 |
| *C. elegans: spo-11(me44)/mls11 IV; bcls39 (ced-1::GFP) V* | Hayashi et al. (10);<br>Stamper et al. (11) | CA376 |
| *C. elegans*: *ojls9[Ppie-1::zyg-12::gfp],unc-24/nT1[qls50] IV;sun-1(jf18)/nT1[qls50] V* | Sato et al. (12); | CA444 |
| *C. elegans*: *spo-11(me44)/nT1[qls50] IV; sun-1(jf18)/nT1[qls50] V* | This paper | CA1788 |
| *C. elegans*: *spo-11(me44)/nT1[qls50] IV; bcls39 (ced-1::GFP)/nT1[qls50] V* | This paper | CA1789 |
| *C. elegans*: *kim27[plk-2::mRuby] I* | Brandt et al. (13); | CA1274 |
| *C. elegans*: *kim27[plk-2::mRuby] I; pch-2(tm1458) II;* | This paper | CA1790 |

**Table S3.**
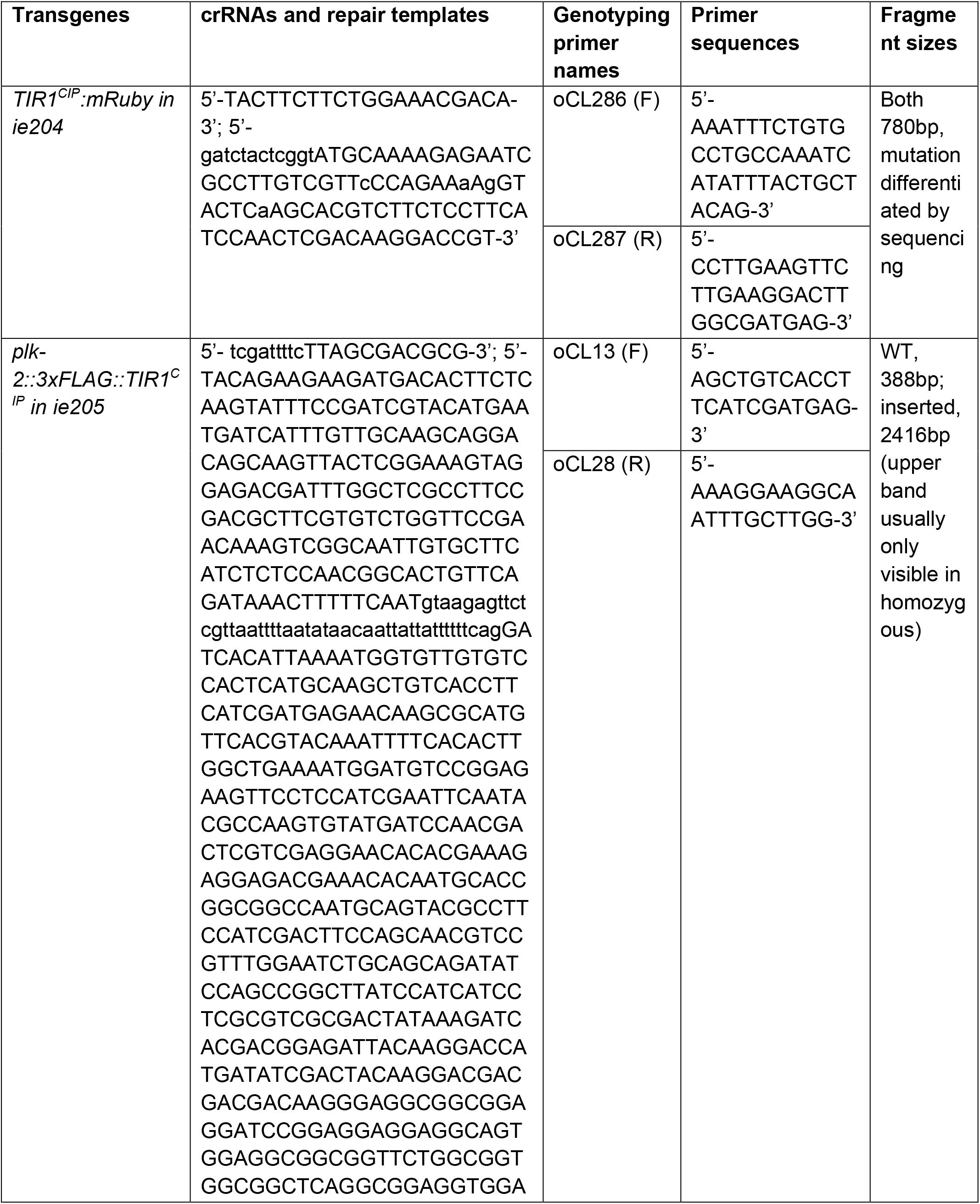
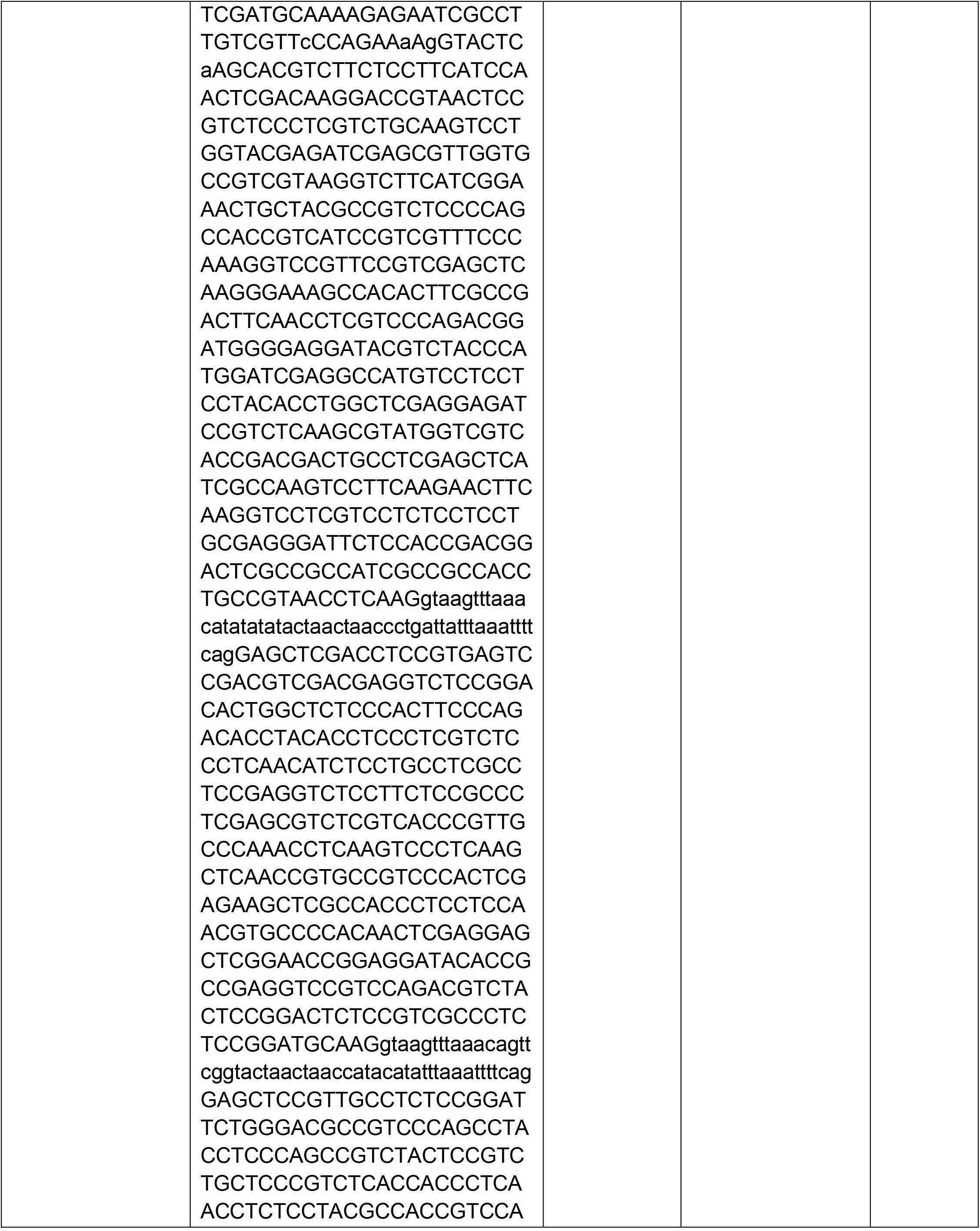
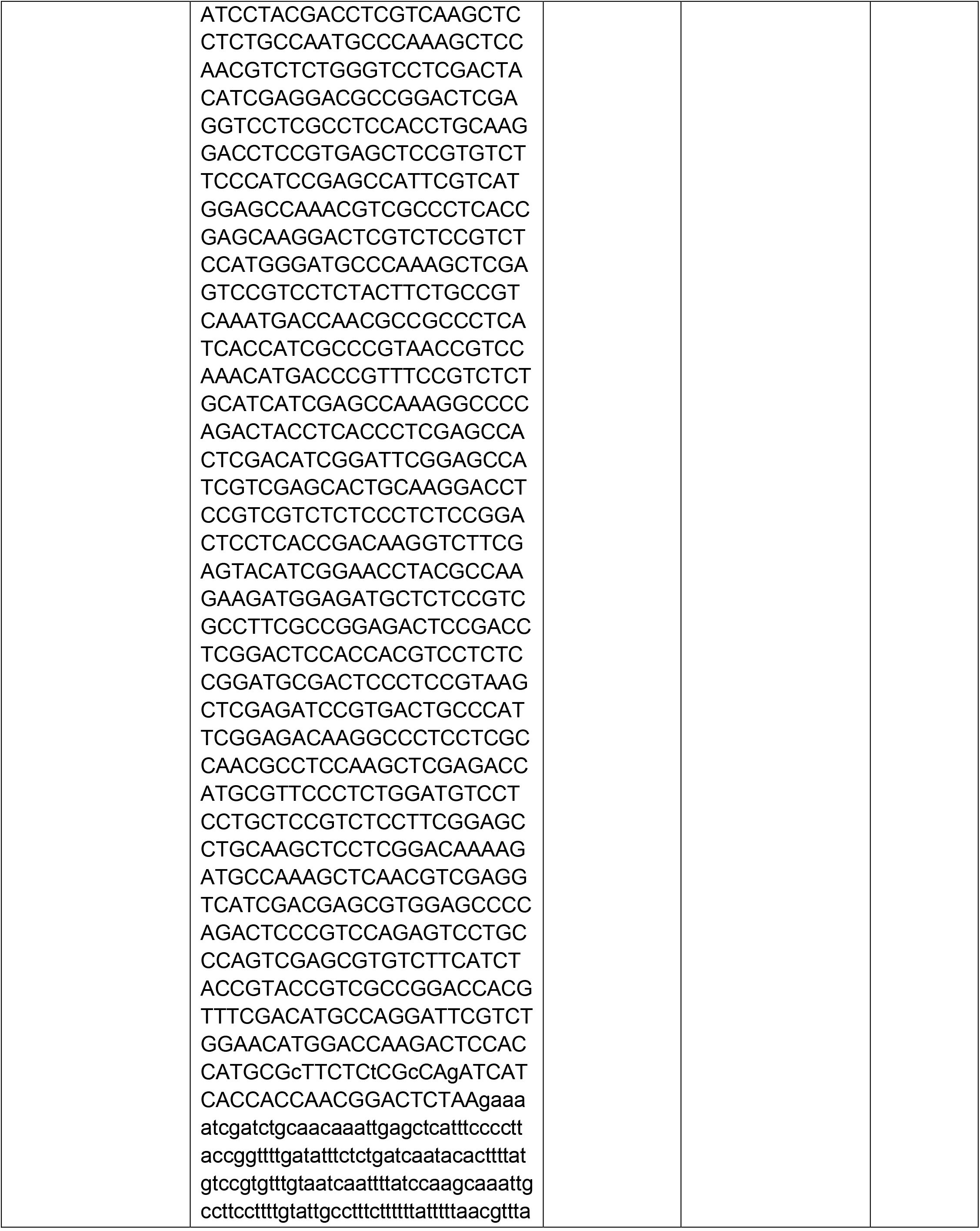
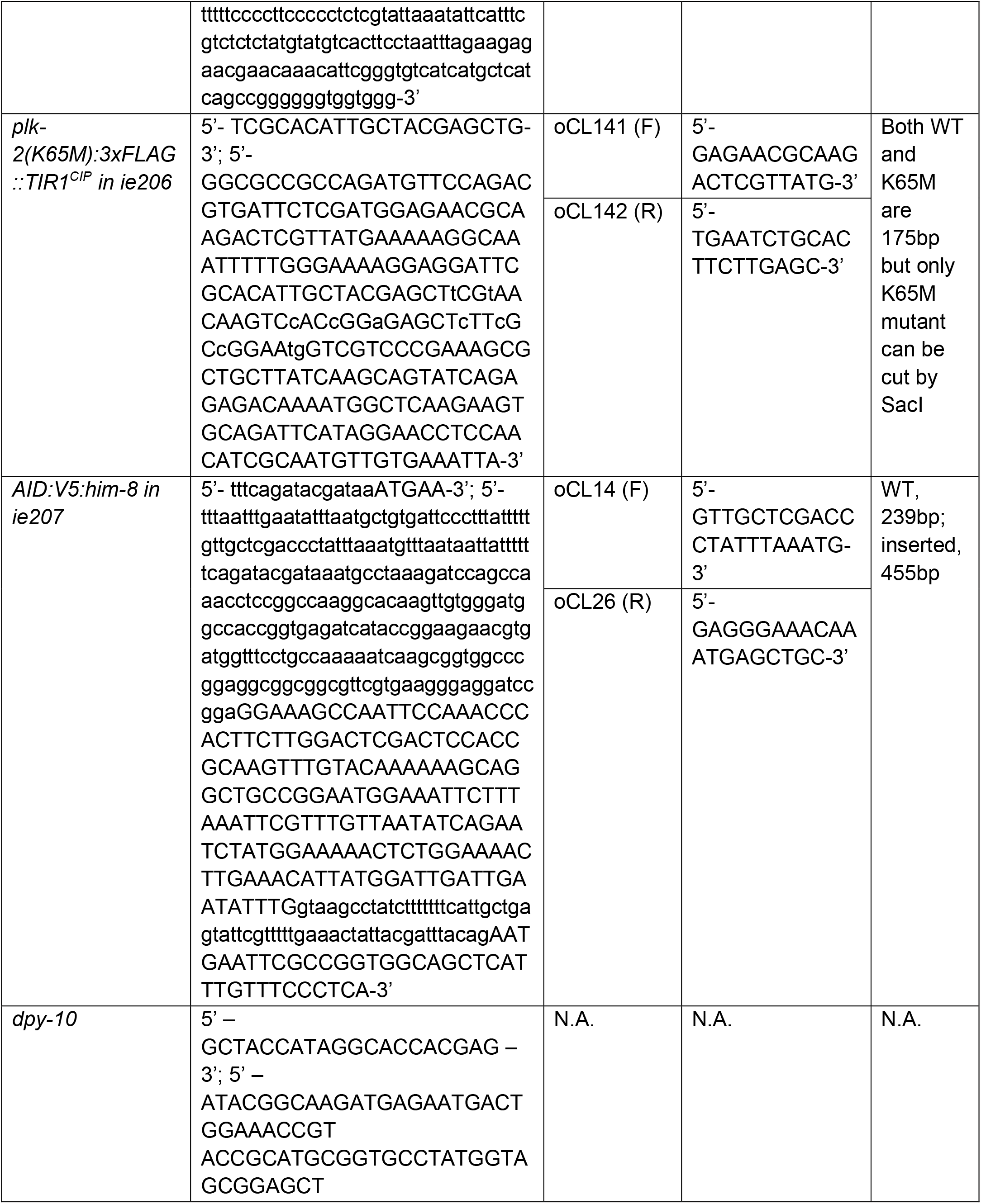
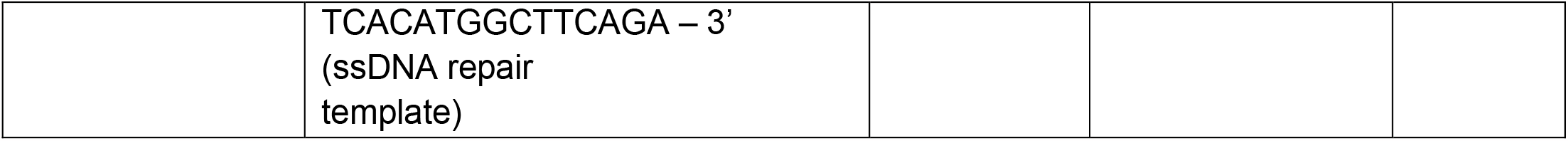
Sequences of crRNAs, repair templates, and DNA primers used to genotype edited progeny.

**Table S4.** RNAi clones used in this study. Both RNAi constructs were confirmed using Nanopore sequencing of whole-plasmids (Primordium).

| Target gene | GenePairs Name | Plate | Well |
| --- | --- | --- | --- |
| *lmn-1* | DY3.2 | 14 | D12 |
| *syp-2* | C24G6.1 | 138 | B6 |

